# The RNA-RNA interactome between a phage and its satellite virus reveals a small RNA differentially regulates gene expression across both genomes

**DOI:** 10.1101/2022.04.08.487710

**Authors:** Drew T. Dunham, Angus Angermeyer, Kimberley D. Seed

## Abstract

Phage satellites exhibit various regulatory mechanisms to manipulate phage gene expression to the benefit of the satellite. While small RNAs (sRNAs) are well documented as regulators of prokaryotic gene expression, they have not been shown to play a regulatory role in satellite-phage conflicts. *Vibrio cholerae* encodes the phage inducible chromosomal island-like element (PLE), a phage satellite, to defend itself against the lytic phage ICP1. Here we use Hi-GRIL-seq to identify a complex RNA-RNA interactome between PLE and ICP1. Both inter- and intragenome RNA interactions were detected, headlined by the PLE-encoded *trans*-acting sRNA, SviR. SviR regulates both PLE and ICP1 gene expression uniquely, decreasing translation of ICP1 targets and affecting PLE mRNAs processing. The striking conservation of SviR across all known PLEs suggests the sRNA is deeply rooted in the PLE-ICP1 conflict and implicates sRNAs as unidentified regulators of phage-satellite interactions.

## Introduction

Conflict between lytic bacteriophages and their hosts imposes tremendous selective pressure on both the virus and host to outcompete each other, resulting in a dynamic co-evolutionary arms race (Koskella and Brockhurst, 2014). To overcome phage predation, bacteria have acquired vast immune systems capable of blocking phages through unique mechanisms (Bernheim and Sorek, 2020). Interactions between the diarrheal pathogen *Vibrio cholerae* and its phages have been proposed to drive seasonality of cholera outbreaks (Faruque et al., 2005). Two decades of surveillance of cholera patient stool samples from Bangladesh has brought to light the specific ongoing conflict between *V. cholerae* and the lytic bacteriophage ICP1 (Angermeyer et al., 2018; Boyd et al., 2021; Seed et al., 2011), providing a tractable model system to interrogate molecular mechanisms employed by both phage and host to antagonize each other.

To defend itself against ICP1 predation, *V. cholerae* encodes a family of anti-phage mobile genetic elements (MGEs) called phage-inducible chromosomal island-like elements (PLEs) (O’Hara et al., 2017). Ten unique PLEs have been identified, each approximately 20 kb in length and encoding a suite of core genes, shared between all PLEs, and non-conserved accessory genes (Angermeyer et al., 2022). All identified PLE(+) *V. cholerae* contain a single PLE, which lays dormant in the *V. cholerae* chromosome until ICP1 infection. Upon infection, the PLE excises from the chromosome (McKitterick and Seed, 2018), replicates to high copy number (Barth et al., 2020a), and deploys multiple anti-ICP1 factors which completely inhibit the production of ICP1 progeny through various mechanisms (Hays and Seed, 2019; LeGault et al., 2022). Unlike many other generalized anti-phage systems, such as restriction modification (Tock and Dryden, 2005) and CRISPR-Cas (Barrangou et al., 2007), PLE-mediated immunity is specific to ICP1, consistent with the relationship between a viral satellite and a virus. Viral satellites are a class of MGEs which specifically parasitize a single virus, leveraging host and viral encoded features to facilitate satellite spread. Accordingly, PLE is spread to neighboring PLE(-) *V. cholerae* in modified ICP1 virions through transduction (Netter et al., 2021), resulting in population-wide immunity and preservation of PLE. Transcriptomic profiling of PLE and ICP1 during infection revealed the ICP1 transcriptome is nearly unperturbed in the presence of PLE, with only the capsid operon being downregulated by a PLE transcriptional repressor CapR (Barth et al., 2020b; Netter et al., 2021). Additionally, RNA-seq identified two intergenic transcripts, a putative PLE *trans*-acting small RNA (sRNA) and an ICP1 long non-coding RNA (lncRNA), to be the two most abundant transcripts late in phage infection (Barth et al., 2020b) (Figure 1A), implying regulatory RNAs may act as post-transcriptional regulators of the PLE-ICP1 conflict.

**Figure 1.**
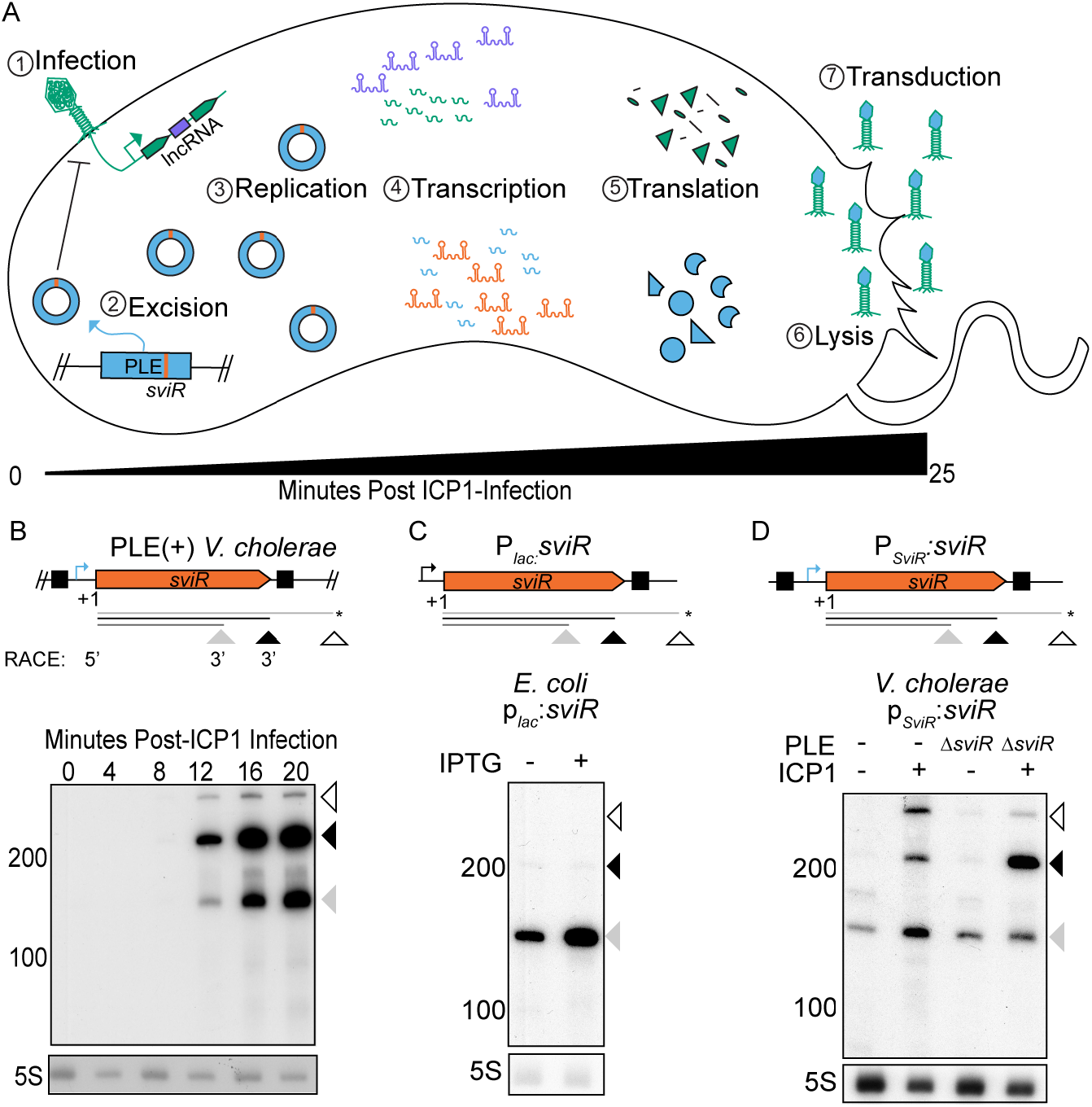
PLE expresses the putative regulatory small RNA SviR in response to ICP1 infection. (A) A representative model of ICP1 phage infection of a PLE(+) *V. cholerae* host. Steps involved in PLE (blue) activation following ICP1 (green) infection are temporally numbered. Upon ICP1 infection, PLE excises from the chromosome (McKitterick and Seed, 2018), replicates to high copy (Barth, Silvas, *et al.,* 2020), and undergoes its transcriptional (Barth, Netter, *et al.,* 2020) and translational programs, resulting in lysis of host cells (Hays and Seed, 2019) and transduction of PLE to neighboring cells in modified ICP1 particles (Netter *et al.,* 2021). ICP1 and PLE regulatory RNAs are shown in purple and orange, respectively. (B) SviR 5’-/3’-RACE products (top, under model) and Northern blot analysis of SviR during a time-course of MOI 2.5 ICP1 infection of PLE(+) *V. cholerae* (bottom). Triangles under gene graphs and on the side of the blots represent the three most abundant species of SviR, shaded according to relative abundance. Annotations below the gene graph represent the SviR TSS, as determined by 5’-RACE and putative stop sites inferred from 3’-RACE data. The species marked with an asterisk correspond to a stop site not confirmed by 3’-RACE. Black boxes on the gene graph represent terminal inverted repeats flanking *sviR.* The native *sviR* promoter prediction is represented by the blue bent arrow icon. (C) Northern blot analysis of ectopic SviR expression from a *p_lac0-1:_sviR* construct in *E. coli.* The P_*lac*_ promoter, represented by the black bent arrow icon, was induced with 1 mM IPTG, expressing the SviR 5’-RACE TSS through the predicted 3’ end of the sRNA. SviR species present in *V. cholerae* are indicated by the arrows present below the gene graph. (D) Northern blot analysis showing ectopic SviR expression from a *p_SviR_:sviR* construct in *V. cholerae.* 100 bp upstream of the SviR TSS was inserted upstream of the same sequence used in (C), providing the predicted SviR promoter sequence (blue bent arrow icon) and left inverted repeat. RNA was isolated from samples grown to an OD_600_ of 0.3, either mock (-) or ICP1 (+) infected at an MOI of 2.5, and incubated for 16 minutes. Host strain PLE genotype indicated above each lane by either (-) for strains lacking PLE or PLEΔ*sviR*.

Bacterial sRNAs are a well-documented class of regulatory RNAs which respond to diverse cellular stresses (Waters and Storz, 2009). sRNA base pairing is initiated with a short seed region of imperfect complementarity to the target transcript (Bandyra et al., 2012) an interaction often, but not always, facilitated by RNA chaperones, such as Hfq and ProQ (Melamed et al., 2020). The result of sRNA regulation varies considerably, with a single sRNA often regulating multiple targets with different outcomes (Feng et al., 2015). The most frequently documented outcome for sRNA regulation is translational inhibition by blocking ribosome association with target transcripts (Storz et al., 2012). However, many unique outcomes of sRNA regulation have been documented and novel mechanisms surely remain to be discovered.

While many sRNAs have been shown to regulate various processes in *V. cholerae* (Hammer and Bassler, 2007; Peschek et al., 2020; Richard et al., 2010), less is known about sRNA regulation between MGEs, phages, and their hosts (Altuvia et al., 2018; Fröhlich and Papenfort, 2016). Both MGEs and prophages have been shown to encode *cis-* and *trans*-acting sRNAs, which regulate transcripts encoded on the core host genome as well as on the MGEs themselves (Bloch et al., 2021; Tree et al., 2014; Wachter et al., 2018). However, MGE- or host-encoded sRNAs regulating lytic phage transcripts have only been predicted from transcriptomic data sets and not yet experimentally validated (Chevallereau et al., 2016). Despite impressive characterization of other phage satellites, such as the P2-P4 system in *Escherichia coli* (Lindqvist et al., 1993) and the *Staphylococcus aureus* pathogenicity islands (Penadés and Christie, 2015), sRNAs have not yet been identified as regulators in satellite-phage conflict.

Recently, multiple techniques to identify targets of bacterial sRNAs *in vivo* have emerged (Han et al., 2016; Melamed et al., 2016; Tree et al., 2014; Waters et al., 2017), each unique in its approach and limitations. Here, we apply High-throughput <u>G</u>lobal s<u>R</u>NA target <u>I</u>dentification by <u>L</u>igation and sequencing (Hi-GRIL-seq) (Zhang et al., 2017), to investigate the role of PLE and ICP1 regulatory RNAs during phage infection of *V. cholerae* under native expression conditions. In doing so, we generated the most thorough view to date of the RNA-RNA interactome between a lytic phage and a host defense element, capturing interactions between phage and satellite transcripts. Characterization of the PLE-encoded sRNA, SviR, identified the regulon of the sRNA, which includes both PLE and ICP1 target transcripts. SviR’s differential regulation of PLE and ICP1 targets positions this sRNA as a key regulator in the PLE-ICP1 conflict. Our findings suggest SviR-target regulation is a conserved feature of PLEs’ manipulation of ICP1, unveiling a complex RNA-RNA landscape between a phage and its satellite.

## Results

### Induction of the PLE regulatory RNA, SviR, is reliant on both PLE and ICP1 factors

To begin characterization of the <u>s</u>atellite-encoded <u>v</u>iral <u>i</u>nduced s<u>R</u>NA (SviR), we first set out to confirm previous transcriptomic predictions that PLE encodes an sRNA (Barth et al., 2020b). RNA-seq suggested SviR is expressed from a PLE intergenic region which contains no predicted open reading frames (ORFs), supporting that the transcript could function as a regulatory sRNA. For all experiments herein we chose to use a co-isolated pair of PLE1(+) *V. cholerae* and ICP1 2011_Dha_A phage (Boyd et al., 2021; Seed et al., 2013), referred to simply as PLE and ICP1. Northern blot analysis of SviR showed the sRNA is robustly detected 12 minutes following ICP1 infection and continues to increase for the duration of the approximately 25 minute ICP1 infection cycle of PLE(+) *V. cholerae* (Figure 1B). SviR expression peaking late in ICP1 infection is consistent with previous RNA-seq data (Barth et al., 2020b) and supports induction of the putative sRNA in response to ICP1 infection.

Northern blot analysis revealed multiple SviR species present during ICP1 infection (Figure 1B). To determine if a single promoter drives expression of all SviR species and identify SviR termination sites, we used a combination of 5’- and 3’-RACE. 5’-RACE confirmed a single 5’ transcriptional start site (TSS) for all sequenced transformants, consistent with the abrupt increase in transcription at that position as observed by RNA-seq (Barth et al., 2020b). Conversely, transformants of 3’-RACE products showed two unique 3’ termination products, one consistent with the shorter 160 nucleotide (nt) SviR species, and another consistent with the longer and more abundant 210 nt SviR species (Figure 1B, below model). No transformants contained a 3’ termination product corresponding to the longer 300 nt species detected by Northern blot, consistent with the low relative abundance of this species compared to the two dominant species. The 3’ termination product corresponding to the most abundant 210 nt species occurs shortly after a predicted rho-independent terminator, likely responsible for transcript termination. No predicted terminator was found corresponding to the shorter or longer species, from which we infer these species are a result of post-transcriptional maturation of SviR. We conclude that expression of SviR is driven from a single promoter upstream of the 5’ TSS which is activated in response to ICP1 infection.

Having determined the bounds of the SviR gene, we next generated plasmid-based constructs aiming to recapitulate expression of the SviR species observed during ICP1 infection of PLE(+) *V. cholerae.* First, we inserted the predicted SviR gene, informed by 5’ and 3’ RACE, into a well characterized vector used for expressing sRNAs in *E. coli* (Corcoran et al., 2012). Induction of this construct in *E. coli* resulted in aberrant SviR expression (Figure 1C) compared to ICP1 infected PLE(+) *V. cholerae* (Figure 1B). While the minor 160 nt SviR species was present, the predominant 210 nt species was nearly undetectable and the larger 300 nt species was not detected. Induction of the promoter did not increase expression of the 210 nt product, suggesting the necessary factors for maturation of SviR were not present in *E. coli* and are likely PLE, ICP1, or *V. cholerae* encoded.

To gain insight into the factors necessary for SviR expression, we next generated a similar plasmid-based construct in *V. cholerae.* In the context of PLE, SviR is flanked by inverted repeat sequences, with the left inverted repeat (IRL) upstream of the SviR TSS and the right repeat (IRR) after the predicted rho-independent terminator. We hypothesized that these inverted repeat sequences and region upstream of the SviR TSS may be involved in SviR expression and maturation. To this end, we inserted the SviR gene and the 100 base pairs (bp) upstream from the SviR TSS into a promoterless plasmid and evaluated SviR expression during ICP1 infection (Figure 1D). Expression of SviR from this construct was induced by ICP1 infection, suggesting the −100 sequence of SviR encodes the native p_*sviR*_ promoter, which is likely directly induced by an unknown ICP1 factor. While it has been known that PLE is transcriptionally activated upon ICP1 infection, this finding provides evidence of ICP1 factors directly activating a PLE promoter.

While p_*sviR*_ expression was increased by ICP1 infection, the relative abundance of SviR species when infecting a PLE(-) host did not match the relative intensity of SviR species observed during a PLE(+) infection. However, introduction of the *p_sviR_:sviR* construct in a PLE(+)Δ*sviR* background followed by ICP1 infection fully recapitulated the length and relative intensity of SviR species as is observed in PLE(+) *V. cholerae* (Figure 1D). We hypothesize the decreased expression of the larger 300 nt SviR species and increased level of the 210 nt species observed in the presence of PLE indicate transcript maturation, potentially due to RNase-mediated processing. Alternatively, it is possible interaction between SviR and PLE target transcripts could lead to transcript turnover, resulting in the observed band patterns. Regardless, proper expression and processing of SviR in *V. cholerae* is dependent on ICP1 and PLE, supporting SviR as a dedicated regulator responsive to ICP1 infection.

To begin to address the role of SviR in PLE-ICP1 dynamics, we evaluated the response of PLE*ΔsviR* to ICP1 infection. The deletion of *sviR* did not grossly impair PLE’s capacity to block ICP1 progeny production, consistent with previous observations that none of the 27 PLE ORFs are necessary for PLE’s inhibition of ICP1 (Hays and Seed, 2019). We detected decreased genome replication of both PLE and ICP1 and a delay in the timing of cell lysis when comparing PLEΔ*sviR* infection to PLE(+) infection (Figure S1). However, both lysis (Hays and Seed, 2019) and genome replication (Barth et al., 2020a; LeGault et al., 2022) are dependent on multiple PLE or ICP1 products, providing no clear candidate genes as targets of SviR regulation. We next pursued identifying target transcripts of SviR during ICP1 infection of PLE(+) *V. cholerae*.

### Hi-GRIL-seq reveals a complex RNA-RNA interactome between PLE and ICP1

To identify putative target transcripts regulated by SviR during ICP1 infection, we turned to Hi-GRIL-Seq (Zhang et al., 2017) to capture RNA-RNA interactions 16 minutes post-ICP1 infection, during peak SviR expression. RNA-seq of T4 RNA ligase treated samples and subsequent bioinformatic analysis (See Methods for full pipeline) identified chimeric transcripts late in ICP1 infection of PLE(+) cells (Figure 2A). PLE-ICP1 cross-genome chimeric transcripts were identified as any transcript where the ends mapped to these two distinct genomes. For reads mapping to a single genome, transcripts were assumed to be chimeric only if the two ends of the reads mapped greater than 1 kb apart, comparable to prior applications of Hi-GRIL-seq (Zhang et al., 2017). This approach identified consistent numbers of chimeras between replicates (Figure S2).

**Figure 2.**
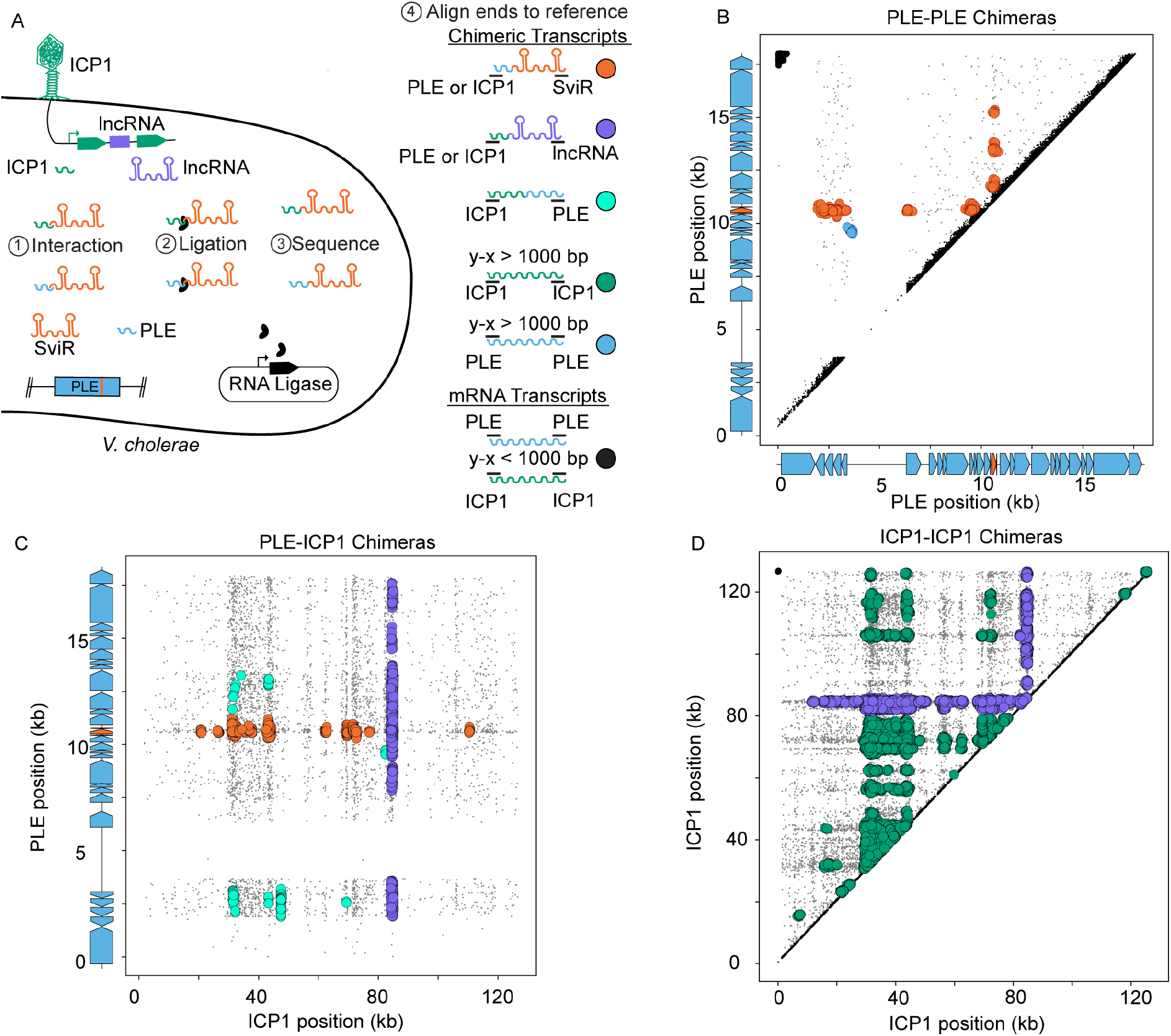
Hi-GRIL-seq during ICP1 infection detects a complex RNA-RNA interactome between PLE and ICP1 transcripts. (A) Schematic of Hi-GRIL-seq carried out during ICP1 infection of PLE(+) *V. cholerae.* Different species of transcript, both chimeric (colors) and non-chimeric (black), are indicated to the right of the model. Assigning the genomic coordinates of the first and last 25 bp of each transcript provides an x-y pairing which was graphed for each read. Reads which map to the same genome closaer than 1 kb between ends were treated as background reads (black), likely representing a single transcript. (B, C, and D) The first and last 25 bp of Hi-GRIL-seq reads were mapped to the reference genomes of PLE-PLE transcripts (B), PLE-ICP1 transcripts (C), and ICP1-ICP1 transcripts (D). For PLE axes, gene graphs on the graph axis demarcate protein encoding ORFs in blue and *sviR* in orange. For PLE-PLE (B) and ICP1-ICP1 (D) interactions, upper triangle graphs were used limit redundant mapping. Reads were clustered using DBScan (Schubert *et al.,* 2017) where greater than 20 reads mapping to an area of diameter 250 bp (B), 350 bp (C), and 1000 bp (D), to scale for differences in genome size between PLE and ICP1, are indicated as a circle. Clusters of chimeras were colored according to the legend in (A). Reads not passing the threshold of clustering were colored gray and sized smaller than reads falling into a cluster.

First, we analyzed the chimeric reads formed between PLE-PLE transcripts (n=1,289 ±196). After DBScan clustering (Ester et al., 1996) to separate background noise from clusters of interactions, all but one cluster of PLE-PLE chimeric reads corresponded to a SviR-PLE suggesting SviR regulates targets across multiple PLE operons (Figure 2B). The majority of these interactions were found to occur between SviR and the 5’-UTR of target transcripts, consistent with canonical sRNA regulation. To our surprise, the only cluster of PLE-PLE reads which was not a SviR-PLE interaction occurred between the 5’-UTR of the operon for PLE *orfs2-5* and PLE *orf12* (Figure 2B), supporting this 5’-UTR as a potential second regulatory RNA encoded by PLE. RNA-seq previously identified this 5’-UTR as being transcribed during ICP1 infection (Barth et al., 2020b). The same promoter responsible for driving SviR expression is conserved upstream of this putative regulatory RNA, suggesting both SviR and the *orfs2-5* 5’-UTR transcript may be expressed by the same factors in response to ICP1 infection. Both of the two putative regulatory RNAs were found by Hi-GRIL-seq to interact with the *orfs12-12.1* operon, placing this target operon at the center of the PLE RNA-RNA interactome.

Parallel analysis of chimeric transcripts between PLE and ICP1 (n=5,399±1,407) revealed a complex interactome between PLE and ICP1 (Figure 2C). Consistent with PLE-PLE chimeras, SviR was the PLE transcript most often ligated with ICP1 transcripts and chimeras were most frequently detected between 5’-UTRs of transcripts and the sRNA. SviR-ICP1 chimeras were detected across multiple ICP1 operons, identifying a list of candidate target genes for SviR regulation of ICP1. The abundance of interactions between SviR and ICP1 targets is consistent with SviR functioning as a cross-genome regulator between ICP1 and PLE. Unlike PLE-PLE chimeras, where only one cluster did not map to SviR, we identified multiple clusters of PLE-ICP1 chimeras which did not include the sRNA. The majority of these non-SviR clusters were formed between PLE transcripts and the putative ICP1-encoded lncRNA, supporting the lncRNA as a potential base pairing RNA encoded by ICP1. In agreement with this, we also observed ICP1-lncRNA chimeras above the detection threshold across ICP1’s genome, showing RNA-RNA interactions between the lncRNA and both PLE and ICP1 transcripts (Figure 2D). Northern blot analysis of the lncRNA showed a similar expression pattern to SviR (Figure S3), showing the lncRNA is robustly transcribed late in ICP1 infection. The lncRNA is a conserved feature of all 67 ICP1 isolates sequenced to date, supporting a role in ICP1 fitness (Boyd et al., 2021). While neither the mechanism of non-specific ligation nor the outcome of the lncRNA-transcript interactions have been investigated in this study, these data indicate that uncharacterized bacteriophage lncRNAs may play a role as regulatory RNAs in phage-MGE conflicts.

### SviR interacts with PLE and ICP1 transcripts with unique seed regions

The RNA-RNA chimeras detected between SviR and both PLE and ICP1 transcripts strongly support SviR functioning as a base pairing sRNA. We next set out to identify the seed region that SviR uses to initiate base pairing with PLE and ICP1 transcripts. Using IntaRNA (Busch et al., 2008; Mann et al., 2017; Raden et al., 2018), we predicted interactions between SviR and the −75 to +25 sequence for every ORF in both PLE and ICP1. From this analysis, we identified two putative SviR seed regions; one interacting with ICP1 targets (Figure 3A, Table 1) and a second interacting with PLE targets (Figure 3B, Table 1). Both seed regions were predicted to interact with target transcripts near the 5’-UTR, consistent with canonical sRNA regulation.

**Figure 3.**
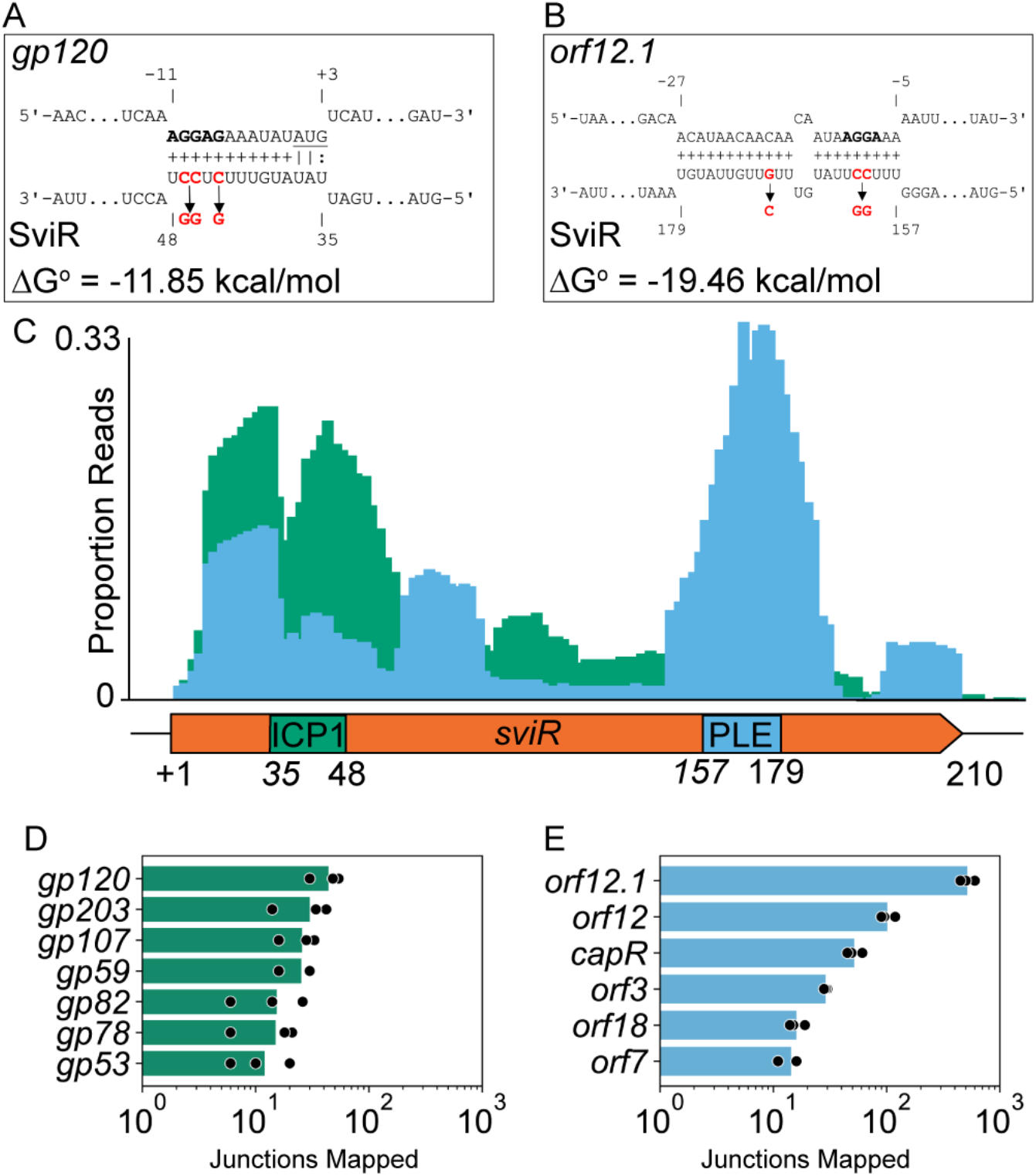
SviR has two unique seed regions to regulate ICP1 and PLE transcripts. (A and B) IntaRNA predicted interactions between the most abundant SviR targets identified by Hi-GRIL-seq in ICP1, *gp120* (A), and PLE, *orf12.1* (B). Each predicted ribosome binding site is bolded and the *gp120* start codon is underlined. Numbering on target transcripts is relative to the start codon and numbering on SviR is relative to the SviR 5’ TSS as determined by 5’ RACE. Nucleotides highlighted in red are predicted by CopomuS (Raden *et al.,* 2020) as important for base pairing and were mutated for subsequent analysis using SviR^ICP1^* and SviR^PLE^* alleles. (C) The first 25 SviR bp flanking the SviR-target junction were mapped to SviR for all SviR-PLE (blue) and SviR-ICP1 (green) chimeras detected by Hi-GRIL-seq. Mapped read junctions across SviR were graphed as a proportion of total SviR-target chimeras in each genome (PLE or ICP1). Annotations on the SviR gene graph below the histogram represent the seed regions predicted by IntaRNA for ICP1 *gp120* (A) and PLE *orf12.1* (B), which are representative of seed regions predicted for most PLE and ICP1 targets (Table 1). (D and E) Quantification of SviR target ORFs in both ICP1 (D) and PLE (E). Mapping of the 25 bp flanking each SviR junction shows the target transcript for each SviR chimera. The maximal peak of reads mapping to a position on the target transcripts is reported for each target, requiring a minimum peak value of 10 reads per target transcript. Reads mapping to intergenic loci were assigned to the downstream *orf* if the mapping was within 50 bp of the *orf* start codon. Presented values are averages of reads mapped across three biological replicates.

**Table 1.** Chimeric junctions mapped from three Hi-GRIL-Seq replicates and IntaRNA interactions. The number of SviR-Target chimeric junctions with an average peak value of at least 10 junctions and the predicted IntaRNA interactions for relevant targets. Junctions mapping to the 5’ −50 sequence relative to the start codon of an ORF were attributed to the downstream ORF. Junctions mapping to targets not predicted to interact with SviR by IntaRNA are qualified with N/A. Target ranges are relative to the start of translation, when applicable, and refer to predicted base pairing interactions as determined by IntaRNA

| Target Transcript | Genome | Hi-GRIL-Seq Chimeras | SviR Range | Predicted Interaction $\Delta G^\circ$ | Target Range |
| --- | --- | --- | --- | --- | --- |
| *orf12.1* | PLE | 516±77.25 | 157 to 179 | -19.46 kcal/mol | -27 to -5 |
| *orf12* | PLE | 101.33±15.5 | 155 to 179 | -10.42 kcal/mol | -28 to -4 |
| *capR*<br>(*orf2*) | PLE | 51.66±8.3 | N/A | N/A | N/A |
| *gp120* | ICP1 | 44±12.5 | 35 to 48 | -11.85 kcal/mol | -11 to +3 |
| *gp203* | ICP1 | 30±14.4 | 37 to 64 | -21.91 kcal/mol | -35 to -2 |
| *orf3* | PLE | 29±1 | N/A | N/A | N/A |
| *gp107* | ICP1 | 25.6±8.7 | N/A | N/A | N/A |
| *gp59* | ICP1 | 25.3±8.1 | 41 to 63 | -17.83 kcal/mol | -20 to +2 |
| *orf18* | PLE | 19±2.64 | N/A | N/A | N/A |
| *gp82* | ICP1 | 15.3±10 | 20 to 48 | -12.86 kcal/mol | -12 to +22 |
| *gp78* | ICP1 | 15±7.9 | N/A | N/A | N/A |
| *orf7* | PLE | 14.33±2.8 | 150 to 177 | -16.85 kcal/mol | -2 to -21 |
| *gp53* | ICP1 | 12±7.2 | 33 to 52 | -23.14 kcal/mol | -7 to -27 |

To support these IntaRNA predictions, we analyzed the junctions of the chimeras from the Hi-GRIL-seq dataset. We reasoned that T4 ligase-mediated junctions between SviR and target transcripts are likely informative for identifying the SviR seed region, as interacting bp are in proximity, which is necessary for ligase activity. For each chimeric read, we identified the first 25 bp of SviR proximal to the chimeric junction and mapped this 25mer back to the SviR coding sequence. Junction reads mapped to SviR primarily in three peaks, indicating three regions of sRNA-target interaction. One peak of chimeric junctions corresponded to the predicted ICP1 seed region, another to the predicted PLE seed region, and a third peak immediately downstream of SviR’s 5’ TSS (Figure 3C). The third peak is likely an artifact, representing non-specific ligation between the 5’ phosphoryl terminus of SviR and target transcripts. Using the same SviR-target junctions, we mapped the target transcript 25mer flanking the junction to its respective genome and quantified the number of chimeras mapping to each target transcript. Both PLE (n=6, Figure 3D) and ICP1 (n=7, Figure 3E) ORFs were predicted to be targets of SviR from this analysis, confirming many IntaRNA target predictions. Many PLE ORFs were predicted by IntaRNA to interact with SviR using the PLE seed region but were not detected by Hi-GRIL-Seq (Table S1). These data identify two sets of candidate genes of SviR regulation and suggest SviR employs two unique seed regions for interacting with PLE and ICP1 transcripts respectively.

### SviR translationally downregulates ICP1 Gp120, a putative ICP1 structural protein

To investigate the outcome of SviR regulation of an ICP1 target, we focused on the most abundant SviR-ICP1 chimera, *gp120* (Figure 4A, Table 1), a gene of unknown function expressed late in ICP1 infection. Hi-GRIL-seq junction mapping detected SviR interactions with the 5’-UTR of *gp120,* just downstream of the TSS identified by 5’-RACE (Figure 4B). Northern blot analysis of *gp120* during ICP1 infection showed equivalent transcript levels in the presence and absence of SviR (Figure 4C). Because SviR is predicted to occlude the *gp120* ribosome binding site (RBS), potentially inhibiting translation, we next engineered a C-terminal 3x-FLAG tagged *gp120* in the native locus and monitored Gp120::3x-FLAG levels during phage infection by Western blot analysis. Gp120 protein levels increased in the PLEΔ*sviR* background (Figure 4D), consistent with SviR regulating Gp120 expression post-transcriptionally by occluding the RBS and blocking translation. Complementation of SviR from *P_SviR_:sviR* decreased Gp120::3x-FLAG levels, confirming the role of SviR in downregulating of Gp120. To assess the role of the SviR seed regions in Gp120 downregulation, we introduced mutations to disrupt each seed region individually (Figures 3A and 3B), as informed by CopomuS (Raden et al., 2020). Surprisingly, complementation with either *P_SviR_:sviR^ICP1*^* or *P_SviR_:sviR^PLE*^* phenocopied PLEΔ*sviR* Gp120 levels*. gp120* base pairing interactions are only predicted with SviR’s ICP1 seed region and we found no evidence of interaction between *gp120* and the PLE seed region by IntaRNA or Hi-GRIL-seq. Both mutant seed regions were expressed to comparable levels to the wild-type allele (Figure S5). It is possible that recruitment of other factors responsible for translational downregulation requires the PLE seed region, or that aberrant secondary structure of the SviR^PLE^* allele prevents proper base pairing between *gp120* and SviR. Nevertheless, these data show SviR-dependent regulation Gp120 translation during ICP1 infection.

**Figure 4.**
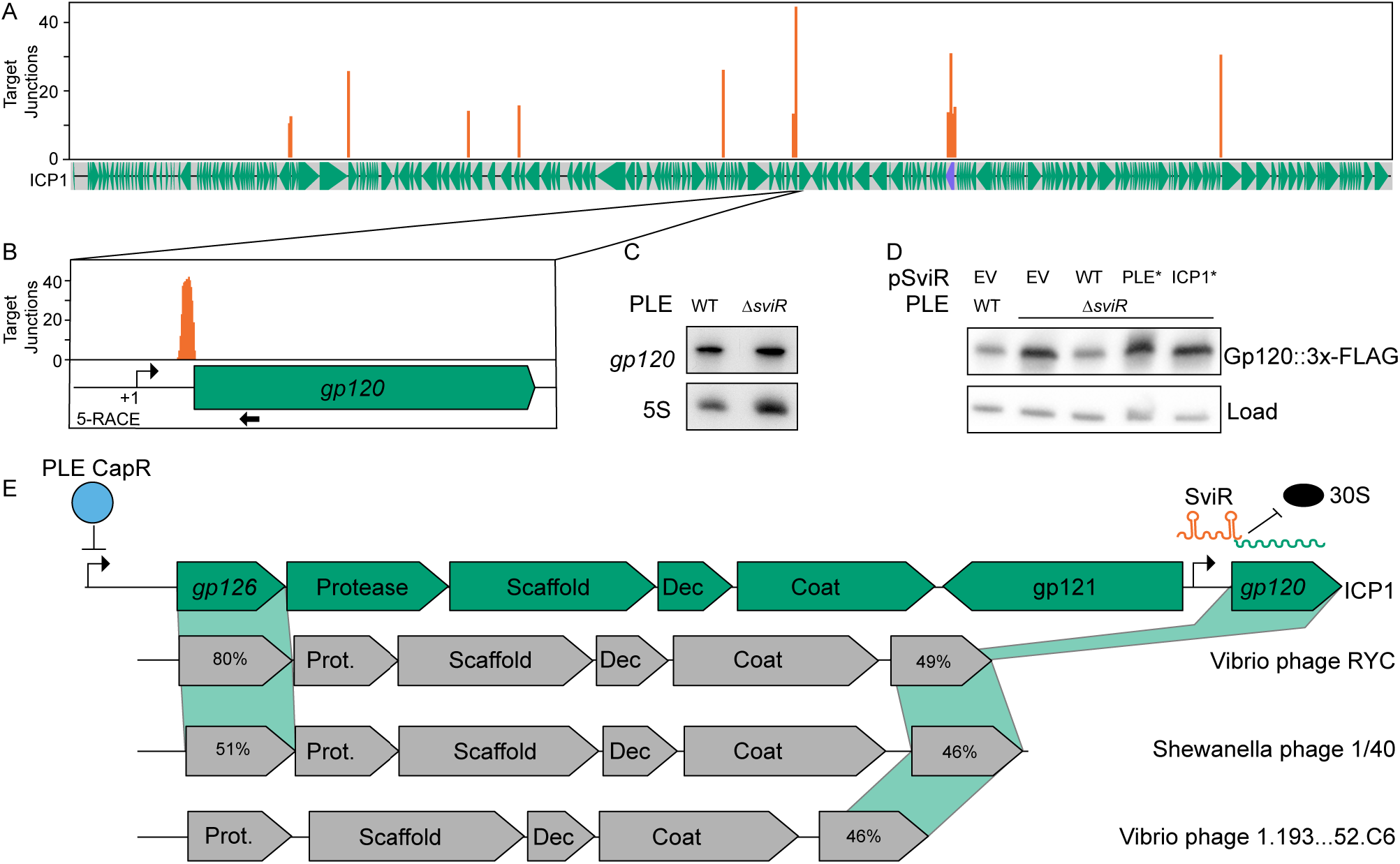
ICP1 Gp120 translation is decreased in the presence of SviR. (A) Mapping of SviR-ICP1 chimeric junctions identified in Figure 3D to the ICP1 genome. The 25 bp following each SviR-ICP1 junction was mapped to the ICP1 reference genome. Mapping represents the average of three biological replicates and positions with an average number of reads less than 10 are not shown. Purple annotation on the ICP1 gene graph represents the ICP1 lncRNA. For individual replicates and unfiltered graphs, see Figure S4. (B) Inset of SviR chimeras mapping to *gp120,* the target with the highest peak of SviR-ICP1 chimeras. Relative position of the *gp120* promoter is represented by a bent arrow, as determined by 5’ RACE. Probe used for *gp120* Northern blot in (C) is indicated by black arrow under the gene graph. (C) Northern blot analysis using a probe complimentary to *gp120.* All RNA samples were isolated 16 minutes post-infection by ICP1, either in a PLE(+) or PLEΔ*sviR* background. (D) Western blot analysis of Gp120::3x-FLAG ICP1 infection in the same host backgrounds and combinations of SviR expression constructs used in (C). p_*SviR*_:SviR^ICP1*^ and p_*SviR*_:SviR^PLE*^ mutant alleles contain mutations highlighted in Figure 3A and 3B, respectively. Total protein input was normalized across each replicate. Load normalization represents a non-specific FLAG background band, present at equal concentrations across samples. (E) Gene graphs of ICP1 and non-related phages showing synteny of the capsid operon and a model of SviR-based regulation of *gp120.* CapR, PLE’s transcriptional repressor, binds upstream of *gp126,* downregulating expression of ICP1’s capsid operon (Netter *et al.,* 2021). *gp121* encodes a putative mobile homing endonuclease gene, hypothesized to have been acquired to de-couple *gp120* from CapR-mediated repression of the capsid operon. However, SviR regulation restores PLE downregulation of *gp120.* Functional annotations of related phages are based on PFam domain predictions. For gene products without annotation, percent amino acid similarity to ICP1 Gp126 or Gp120 was shown instead, as calculated by EMBOSS Needle alignment. Dec = Decoration protein; Prot. = scaffold protease.

Notably, Gp120 downregulation by SviR represents the second PLE-encoded mechanism of downregulating this region in ICP1, which includes the capsid operon. PLE-encoded CapR is a transcriptional repressor that binds to the promoter region of the capsid operon upstream of *gp126* (Figure 4E) (Netter et al., 2021). However, *gp120* is transcriptionally de-coupled from the ICP1 capsid operon, due to *gp121* encoded on the reverse strand. *gp121* is a putative homing endonuclease gene, a type of selfish MGE hypothesized to play a role in shaping PLE-ICP1 coevolution (Barth et al., 2021; Boyd et al., 2021; Netter et al., 2021). As CapR can no longer transcriptionally repress *gp120* in this context, SviR downregulation of Gp120 translation may act to restore the absent CapR downregulation. In support of this model, multiple phages lack a *gp121* homolog but otherwise share gene synteny with ICP1’s capsid operon (Figure 4E). PSI-BLAST of these *gp120* homologs detects similarity to the T4 *inh* protein, an inhibitor of the procapsid protease in coliphage T4, supporting *gp120* may play a role in ICP1 capsid morphogenesis (Miller et al., 2003). This adaptation-counter adaption model provides an example of the diverse evolutionary mechanisms employed in response to phage-MGE conflict.

### Transcripts from multiple PLE operons are regulated by SviR

Next, we set out to determine the outcome of SviR-based regulation of PLE targets. Mapping of junctions across PLE revealed interactions specifically with the 5’-UTR of target sequences across multiple PLE operons (Figure 5A). To characterize the outcome of these interactions, we focused on the most abundant chimeras detected by Hi-GRIL-Seq, those between SviR and *orf12.1* (n=516) and *orf12* (n=101) (Figure 5B, right). PLE *orf12* and *orf12.1* are both small *ORFs* of 59 and 39 amino acids, respectively, each encoding proteins of unknown function with homologs only in other PLEs. We used Northern blot analysis with a probe complimentary to *orf12.1* to assess the fate of the operon transcript including *orf12* and *orf12.1.* In the presence of SviR, shorter transcript bands, consistent with degradation products, were observed for *orf12.1* (Figure 5C), which were no longer present in PLEΔ*sviR.* Complementation of SviR from p_SviR_:*sviR* restored the shorter transcripts, showing SviR regulation is necessary for regulation of this operon. Importantly, complementation was successful with p_SviR_:*SviR*^ICP1*^, but not p_SviR_:*sviR*^PLE*^, supporting the importance of base pairing of SviR’s PLE seed region for wild-type regulation. The same transcript pattern was observed using a probe against *orf12,* showing the outcome of transcript regulation is consistent across the operon (Figure S5A). In support of direct regulation of *orf12.1* by SviR, expression of an *orf12.1::GFP* translational fusion reporter was decreased in the presence of SviR in a PLE seed region dependent manner (Figure 5D, Figure S6B). However, introduction of compensatory mutations to restore interaction between SviR^PLE*^ and the *orf12.1* 5’-UTR necessitated mutation of the RBS and reduced GFP expression significantly, precluding us from verifying base pairing using this approach (Figure S6A, S6C).

**Figure 5.**
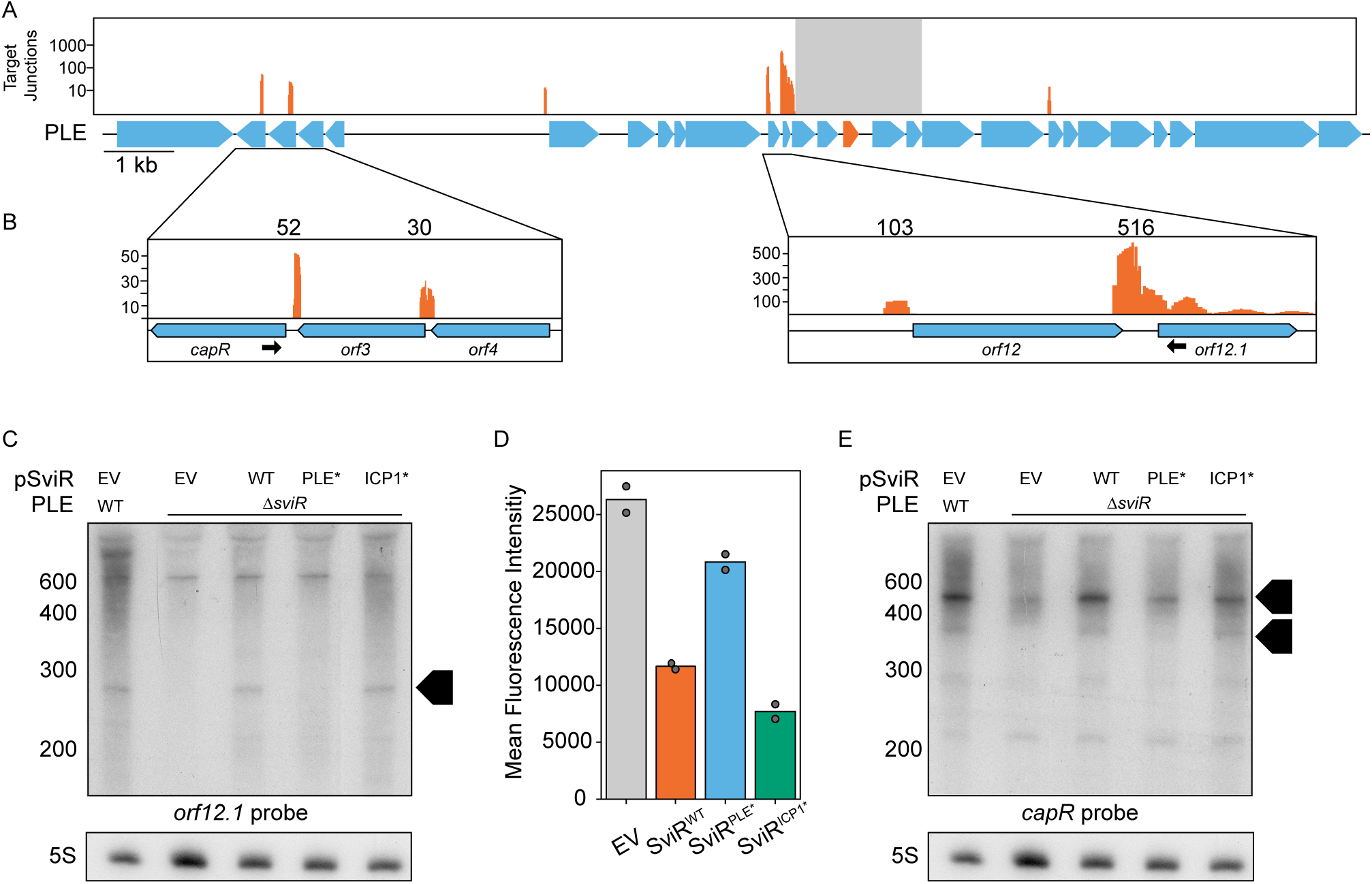
SviR regulation leads to altered levels of transcripts across PLE. (A) Mapping of SviR-PLE chimeras to the PLE reference genome. Mapping was performed as in Figure 4A. Reads mapped represent the average of three biological replicates. For individual replicates and unfiltered graphs, see Figure S4. Reads are mapped on a log scale in (A) due to the range of values mapped. The gray box overlaying the SviR annotation (orange) on the PLE gene graph represents the 1kb region flanking the center of SviR, where chimeric reads were unable to be discerned from nonchimeric transcripts due to proximity to the sRNA. (B) Inset images of PLE target junction mapping. Zoomed insets show linear scale of SviR interactions with *capR* (left) and *orfs12* and *12.1* (right). Black arrows indicate the Northern blot probes used in (C) and (E). Scales between each inset differ based on local maxima. (C and E) Northern blot analysis using a probe complimentary to *orf12.1* (C) or *capR* (E) in the 5’ region of the ORF as indicated in (B). All RNA samples were isolated 16 minutes post-ICP1 infection. SviR^PLE*^ and SviR^ICP1*^ mutant alleles contain mutations highlighted in Figure 3A and 3B. Black boxed arrows highlight transcript species differentially regulated by SviR and dependent on the PLE seed region. Blots were stripped and re-probed with each target probe, resulting in identical 5S images for both *orf12.1* and *capR* blots. (D) Flow cytometry analysis of *orf12.1::GFP* expression in the presence and absence of SviR alleles in *E.coli.* SviR expression was induced in all samples and samples were gated to remove non-fluorescent cells. Graphed values represent the mean fluorescent intensity of the GFP(+) population of 50,000 counted events, analyzed by FlowJo.

A second PLE operon detected by Hi-GRIL-seq to be targeted by SviR was the operon encoding CapR, where both *capR* and *orf3* were detected as targets (Figure 5B, left). To determine if the *capR* operon shared the same fate as the *orf12.1* operon, we performed similar Northern blot analysis using a probe complimentary to *capR.* Consistent with the *orf12.1* operon, we observed differential regulation of *capR* dependent on the PLE seed region of SviR (Figure 5E), suggesting direct interaction between SviR and multiple PLE-encoded target operons. The regulation of target transcripts occurs independent of either RNA chaperone Hfq or ProQ, although Hfq may play a role in transcript abundance of both SviR and targets (Figure S5B-D). It is possible *V. cholerae,* PLE, or ICP1 may encode a novel RNA chaperone, or regulation of these targets occurs independent of chaperones. Nevertheless, differential regulation of multiple PLE targets confirms SviR as a regulator of targets across the phage-MGE boundary, capable of regulating both PLE and ICP1 transcripts.

### SviR regulation is predicted to be a conserved feature of all known PLEs

Alignment of SviR and surrounding sequence from each of the ten PLEs shows high conservation of SviR (Figure 6A). The regions of the highest conservation across the ten SviR alleles are those with functional relevance, including the promoter region, both seed regions, and an intrinsic terminator predicted to terminate SviR transcription to generate the most abundant 210 nt species (Figure 1B). Consistent with these predictions, RNA-seq profiling of PLEs 1-5 during ICP1 infection detected SviR expression consistent with the pattern observed for PLE 1 (Barth et al., 2020b). Because SviR is highly conserved across all PLEs, we next questioned if the SviR targets in other PLEs had predicted base pairing with their cognate SviR allele (Figure 6B). In every PLE, SviR was predicted to interact with the 5’-UTR of targets using the seed region identified here in PLE 1. Some conserved loci, such as the *capR* operon and the cluster of genes downstream of SviR, were consistently predicted to be regulated by SviR across all PLEs. However, the region between PLE *repA* and *sviR* is also predicted to be regulated despite the genetic content of this locus being highly variable. This suggests the role of SviR regulation is central to regulation of PLE gene expression, as SviR targets include core and accessory ORFs in multiple PLE operons.

**Figure 6.**
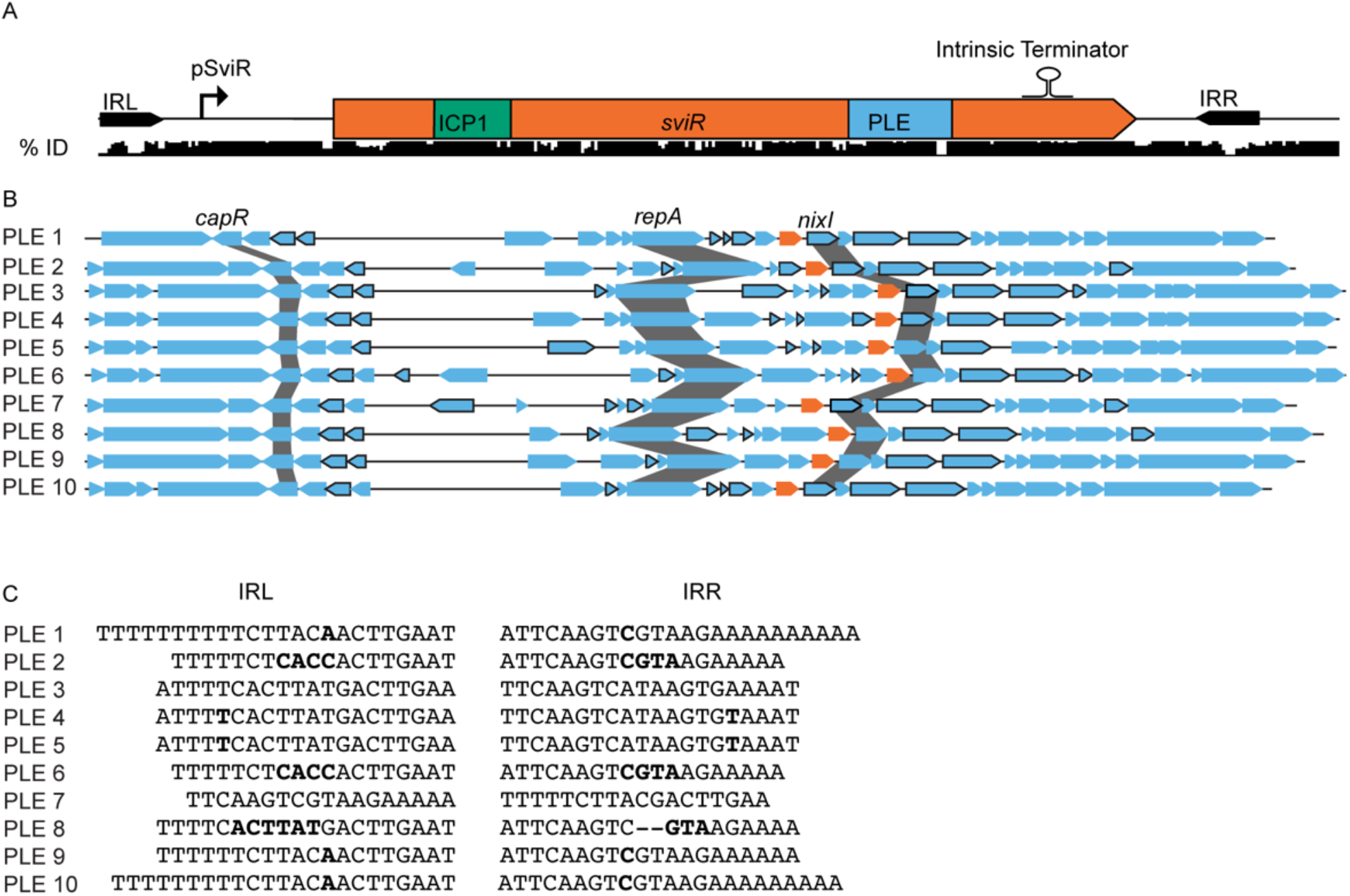
SviR is a core regulatory component of all PLEs discovered to date. (A) SviR and its predicted *cis*-regulatory elements are conserved across all ten PLEs. A nucleotide alignment of SviR and flanking regions (below) showing percent identity of the region across all ten PLEs. The ICP1 and PLE seed region are highlighted in green and blue, respectively. The inverted repeats flanking *sviR* are marked by black arrows and the relative location of the SviR promoter and intrinsic terminator are annotated above the sequence, as determined by RACE. (B) SviR is predicted by conservation and IntaRNA to be a regulatory feature of all ten PLEs. Each of the ten PLEs is shown by a representative gene graph with each SviR allele in orange. ORFs outlined in black are predicted by IntaRNA to interact with the SviR allele encoded in the same PLE. Only SviR-PLE target interactions which were predicted to interact with the PLE seed region with ΔG° < −10 were called putative targets. (C) The inverted repeats flanking SviR from each of the ten PLEs written in the 5’ to 3’ orientation relative to the IRL. Mismatched bases between IRL and IRR are highlighted in bold, and gaps in the repeat without a cognate mismatch are represented by hyphens. Despite the highly variable sequences between the 10 PLEs, the presence of inverted repeats is consistent across all PLEs.

Finally, every SviR allele is flanked by an imperfect inverted repeat sequence, one upstream of the SviR promoter and another following the predicted rho-dependent terminator (Figure 6A). Between the ten PLEs, the sequence of the inverted repeats is variable, but mutations present in one inverted repeat are routinely compensated for in the opposing repeat, suggesting selection for repeat complementarity (Figure 6C). While the role of these repeats is unknown, they are necessary for ectopic expression of SviR (Figure 1D), suggesting the potential role in SviR expression may lead to selection for conservation between repeats. Together, these bioinformatic analyses support SviR as a key factor in PLE-ICP1 co-evolution, regulating PLE and ICP1 targets throughout decades of MGE-phage pairings.

## Discussion

This work describes the role of SviR, a satellite-encoded sRNA, in differential regulation of satellite and phage gene expression during infection. SviR’s function as a regulator of both PLE and ICP1 transcripts places the sRNA at the interface between the satellite and its phage. Satellite parasitism of phages requires a delicate balance of permitting and inhibiting select aspects of phage infection to proceed. Premature or delayed inhibition of phage gene expression is deleterious to the satellite, which requires phage gene products for satellite transmission. To obtain this necessary balance, satellites encode diverse mechanisms to regulate expression of both satellite and phage gene expression. PLEs (Netter et al., 2021), SaPIs (Ram et al., 2014), and P4 (Lindqvist et al., 1993) all downregulate phage late gene expression through unique mechanisms, now including a *trans*-acting sRNA. SviR not only fills this role for PLE, but also regulates PLE targets through a different mechanism, highlighting the versatility of sRNAs as regulators. While PLE gene regulation remains mysterious, characterization of SviR has shown the sRNA is a pivotal piece of the puzzle. Broadly, the RNA-RNA interactome identified in this study brings to question the role regulatory RNAs, which have otherwise not been interrogated, in other phage-satellite conflicts, highlighting the potential for discovery in such systems with global RNA interactome studies.

Given the conservation of SviR among all PLEs, we anticipate SviR regulation to be a consistent pressure faced by ICP1. Notably, all ICP1 targets detected above our Hi-GRIL-seq threshold were core genes, conserved across the 67 ICP1 isolates, supporting SviR regulation targeting essential ICP1 processes. Phages can rapidly overcome anti-phage mechanisms, such as acquisition of the mobile gene Gp121 to disrupt CapR repression. One such potential anti-SviR mechanism encoded by ICP1 may be the presence of a base-pairing lncRNA. The abundance of interactions between the two transcripts (Figure 2C) supports the lncRNA as a potential sponge transcript, acting to sequester SviR from other transcripts during infection and increasing transcript turnover by recruitment of RNases to duplexed RNA (Denham, 2020). Of the few well-characterized lncRNAs in prokaryotes and phages, trans-acting base pairing has not been functionally described (Harris and Breaker, 2018). Another possible explanation for the ICP1 lncRNAs promiscuous base pairing *in trans* is the lncRNA increasing transcript turnover during infection, increasing nucleotide availability for replication and transcription of other phage products. The lncRNA may be central to ICP1’s response to the PLE program and further investigation of the role of lncRNAs in MGE-phage and MGE-host conflict will likely reveal novel lncRNA functions.

The profound conservation of the genomic locus of SviR, being flanked by variable repeat sequences, raises questions about the origins of the sRNA and what pressures select for maintenance of the locus architecture. The conservation of inverted repeats in each PLE with unique sequences between PLEs is of particular interest. Inverted repeats flanking an sRNA are hallmark characteristics of miniature inverted transposable elements (MITEs), a transposase-deficient class of MGE (Wachter et al., 2018). MITE-mediated sRNA mobilization leads to domestication of the MITE in loci where the sRNA is selected for, consistent with SviR integration into PLE. MITE-encoded sRNA maturation can be reliant on inverted repeat sequence (De Gregorio et al., 2005; Mazzone et al., 2001), possibly explaining the conservation of SviR repeat sequences within each PLE. Different transposable elements, the IS200/IS605 family, are also known to mobilize sRNA into diverse loci (Altae-Tran et al., 2021). Interestingly, IS200/IS605 elements are associated with distal nucleases, which are guided by the transposable element-encoded sRNA. The ORF downstream of SviR, NixI, was recently shown to be a nicking endonuclease, implicated in the degradation of ICP1’s genome during rolling circle replication (LeGault et al., 2022). While NixI is not encoded within the transposable element repeats, the upstream sRNA proximal to a nuclease effector raises the possibility of a similar RNA-guided nuclease activity for NixI, secondary to the primary function as a replication inhibitor. Regardless of the evolutionary history of SviR, mobilization of a regulatory RNA into an MGE involved in phage defense is consistent with the “guns-for-hire” model of genetic exchange between hosts and MGEs (Koonin et al., 2020), whereby SviR was domesticated by PLE to assist in the MGE’s anti-phage activity. Collectively, SviR acts as a multi-functional regulator of gene expression in the PLE-ICP1 conflict, revealing a new mechanism of gene regulation to investigate in other MGE-phage conflicts.

## Limitations of this study

Characterization of SviR in the native context of phage infection with Hi-GRIL-seq is a powerful approach, ensuring that target regulation we observed is occurring during natural infection. However, the cell state of ICP1 infected PLE(+) *V. cholerae* introduces countless variables, which we are unable to reconstitute outside of the native context, limiting direct validation of base pairing interactions. Notably, the 25 minute accelerated lifecycle of ICP1 infection of PLE(+) *V. cholerae* is too brief of a timeframe for expression of common fluorophores, such as GFP. Alternative approaches, such as NanoLuciferase (England et al., 2017), were found to be too stable to observe the subtle changes associated with SviR-target regulation.

While we were able to use Hi-GRIL-seq to detect RNA-RNA interactions between multiple loci in both PLE and ICP1, only a single time point of interactions was analyzed. The remaining *V. cholerae* genome may also contain abundant RNA-RNA interactions occurring in response to ICP1 infection. However, during infection, the *V. cholerae* genome is degraded (McKitterick and Seed, 2018), removing the vast majority of non-PLE transcripts late in ICP1 infection (Barth et al., 2020b). A similar workflow applied to early timepoints in the conflict may enrich for other PLE-ICP1 chimeric transcripts, as well as non-PLE *V. cholerae* transcripts involved in response to ICP1 infection.

## Acknowledgments

We thank Dr. Gigi Storz, Dr. Philip Adams, and Dr. Sahar Melamed for their training on Northern blot analysis, general advice, and providing *E. coli* reporter plasmids. We also thank the members of the Seed Lab for their feedback, in particular Dr. Reid Oshiro and Caroline Boyd for help during manuscript revision. D.D. was supported by an NSF Graduate Research Fellowship (Fellow ID no. 2018257700). The project described was supported by Grant Numbers R01AI127652 and R01AI153303 (K.D.S.) from the National Institute of Allergy and Infectious Diseases and its contents are solely the responsibility of the authors and do not necessarily represent the official views of the National Institute of Allergy and Infectious Diseases or NIH. K.D.S. holds an Investigators in the Pathogenesis of Infectious Disease Award from the Burroughs Wellcome Fund.

## Author contributions

Conceptualization, D.T.D. and K.D.S.; Methodology, D.T.D. and K.D.S.; Investigation, D.T.D.; Data Curation, D.T.D. and A.A.; Writing – Original Draft, D.T.D. and K.D.S.; Writing – Review and Editing, D.T.D., A.A. and K.D.S.; Funding Acquisition, D.T.D. (NSF GRFP) and K.D.S.

## Declaration of interests

K.D.S. is a scientific adviser for Nextbiotics Inc. All other authors declare no competing interests.

## Lead Contact

Further requests for resources, strains, and reagents should be directed to and will be fulfilled by the Lead Contact, Kimberley Seed.

## Bacterial and Bacteriophage Culture

### Bacterial

Genotypes of all *Vibrio cholerae and Escherichia coli* strains and plasmids used in this study are listed in the strains and plasmids list. *V. cholerae* strain KDS2 (a clinical isolate harboring PLE1) was used as the wild type strain for genetic manipulations. *E. coli* MG1655 was used as the wild type *E. coli* expression. All overnight bacterial cultures were grown with aeration in a roller at 250 rpm at 37°C in LB medium. Larger volume cultures were grown shaking at 250 rpm at 37°C in LB medium. Ampicillin (100 ug/mL), streptomycin (100 ug/mL), and chloramphenicol (25ug/mL *E. coli,* 2.5 ug/mL *V. cholerae)* were supplemented as necessary. 1 mM IPTG was supplemented to induce cultures when necessary.

### Bacteriophage

All ICP1 listed in this study are derivatives of ICP1 strain 2011_Dha_A, detailed in the strains and plasmids list. Bacteriophage stocks were stored in STE buffer at 4°C. All bacteriophage infections were initiated in a host cell culture grown to OD_600_=0.3, consistent with the onset of logarithmic growth of *V. cholerae.* Infections were either carried out at a multiplicity of infection (MOI) of 2.5 or 0.1, as indicated. Induction of *V. cholerae* expression constructs was performed when cultures reached an OD_600_=0.2 for 20 minutes prior to infection, where MOI calculations were done to final culture OD_600_ value.

## Strains and plasmids

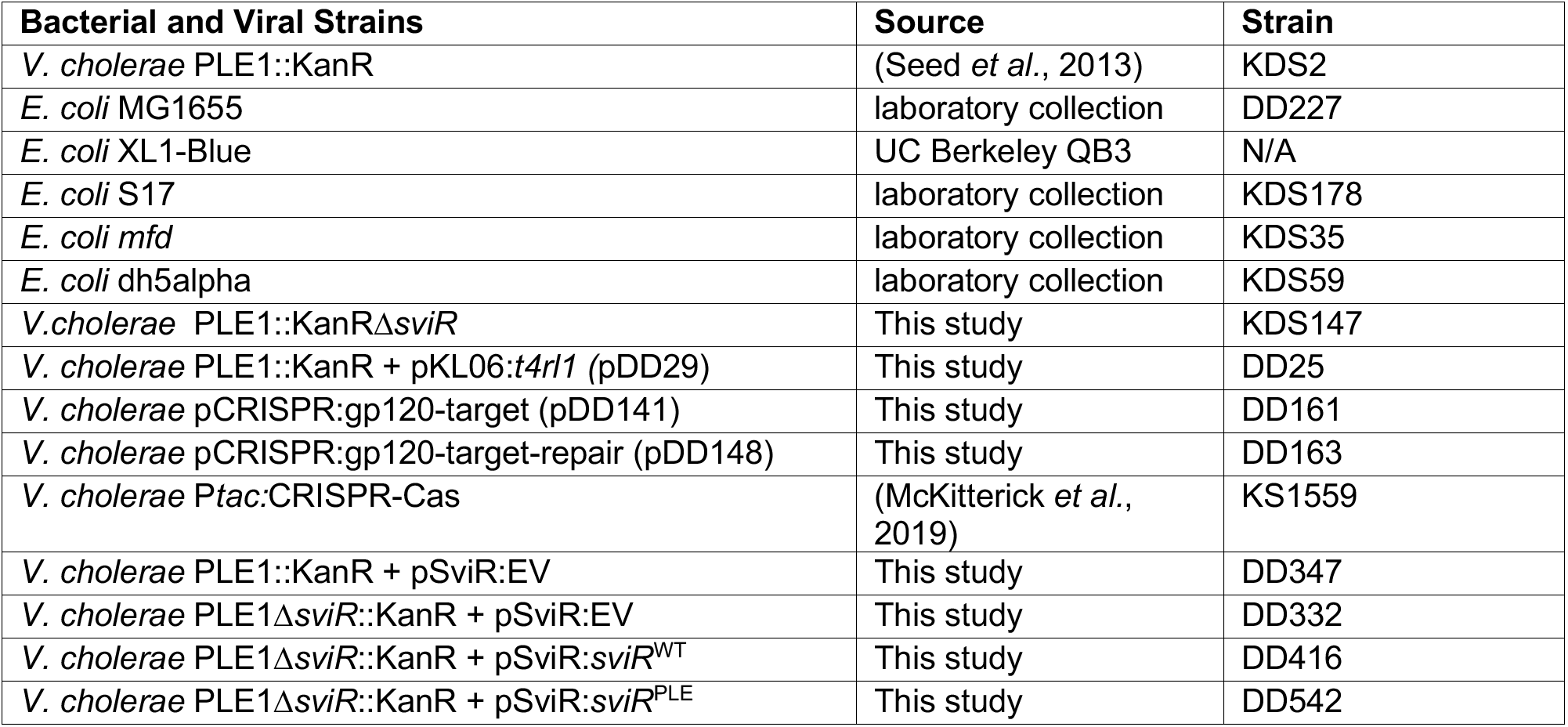
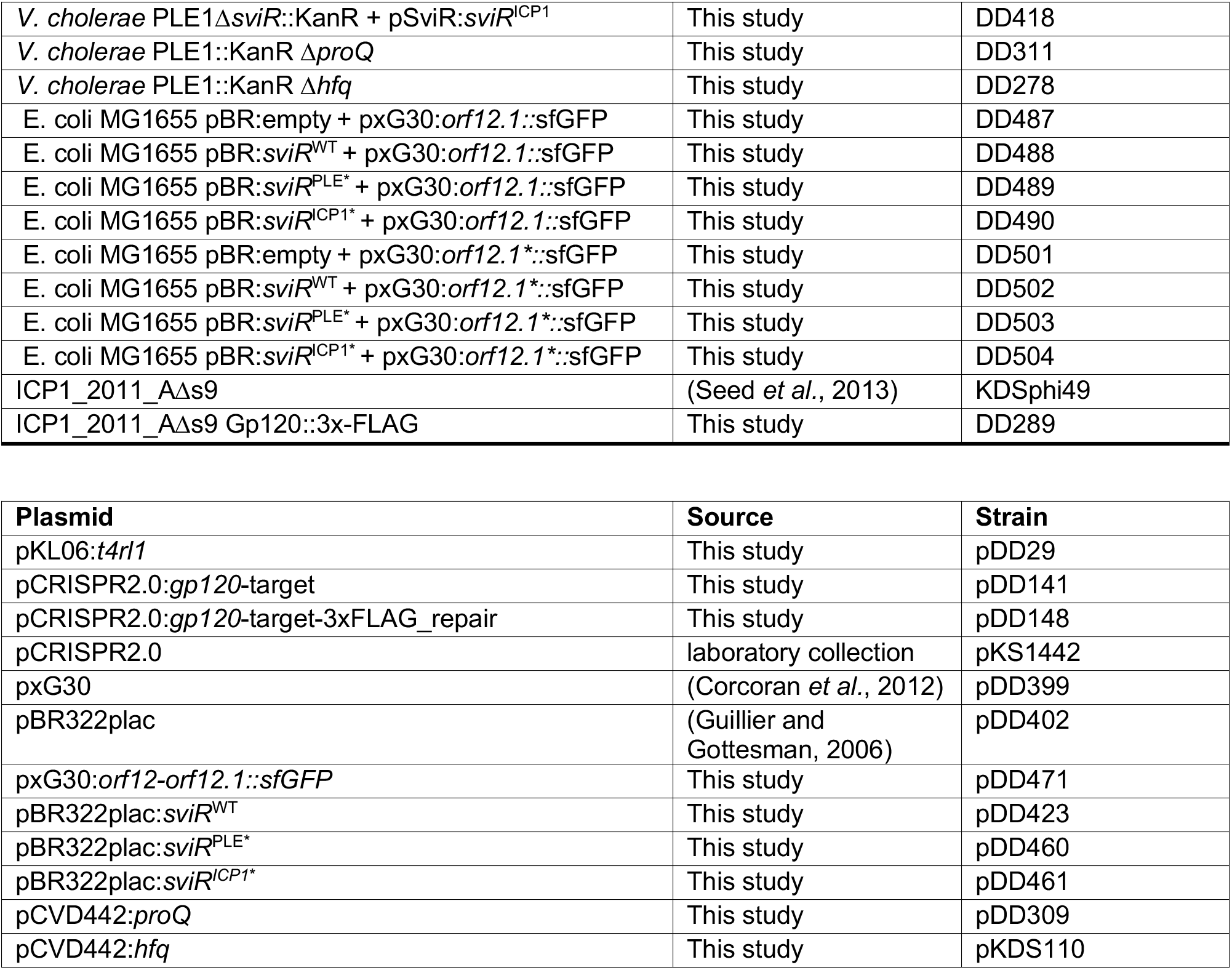

## Materials and Methods

### Hi-GRIL-seq read mapping

Forward and reverse read fastq files were first trimmed with Trimmomatic (Bolger et al., 2014) (leading 30 -trailing 30 -slidingwindow 5:30 -minlen 25) to trim low quality bases from the ends of reads. Trimmed read pairs were merged with PEAR (Zhang et al., 2014) (-j 3 -u 0.1 -p 0.01 -v 10 -n 50), requiring merged read pairs to be a minimum of 50 bp in length with a minimum p-value of 0.01. The first and last 25 bp of each paired read was then mapped to reference genome genbank files for PLE, ICP1, and both *V. cholerae* chromosomes with bowtie2 (Langmead and Salzberg, 2012) (-p 4 -D 1 -R 1 -N 0 -k 1 -I 0 -X 50 --ignore-quals --end-to-end --no-unal). The resulting bam file was parsed with samtools (Li et al., 2009) and any read which mapped to two distinct genomes was considered as chimeric. Reads which mapped to the same genome but had at least 1kb of sequence separating the two ends were also considered chimeric, predicted to be interactions between two transcripts in the same genome. -The terminal mapping positions of all chimeric reads were then recorded as x and y values and plotted on scatter plots against reference genomes. Scatter plots were then analyzed with DBScan (Ester et al., 1996) (-n 10 -eps [min distance]) with the minimum distance between clustered reads varied based on genome size for PLE-PLE (250 bp), PLE-ICP1 (350 bp) and ICP1-ICP1 (1000 bp) chimeras, respectively. Clusters of 10 or more reads where the average x or y value of all reads in the cluster mapped to SviR or the lncRNA were colored according to figure legends. Plots of clustered graphs were generated with matplotlib (Hunter, 2007).

### SviR-chimera junction mapping

Trimmed and merged chimeric reads were aligned to reference genomes using BLASTn (Camacho et al., 2009) default settings. Reads where the top two HSP BLASTn hits mapped to SviR and a target sequence were split at the junction between the alignments and the 25 base pairs closest to the junction site were extracted in fasta format. SviR and target junction fragments were then mapped to respective reference genbank files using bowtie2 with default settings and sorted with samtools. Sorted bam files were used to calculate junction read coverage bedgraphs using bedtools genomecov (Quinlan and Hall, 2010) with default settings.

Mean genome coverage of three replicates was filtered to remove target sites with fewer than 10 reads mapping to a single locus and the filtered mean coverage graphs were visualized with Integrative Genome Viewer (IGV) (Robinson et al., 2011). SviR-target interactions at a given gene of interest were calculated by reporting the peak average coverage of target junctions at a given ORF. Reads mapping to the intergenic region between two transcripts were mapped to the flanking ORFs if the peak was within 50 bp of the gene coding sequence. Values were reported as average ± SEM between replicates.

For SviR junction mapping, SviR-mapping junction coverage was calculated with bedtools genomecov. Coverage across the SviR coding region was normalized to the total number of junctions mapping to SviR in a given replicate, and the average relative frequency of a junction mapping to a given SviR position across three replicates was graphed using IGV.

### IntaRNA and CopomuS sRNA-target predictions

IntaRNA (Mann et al., 2017; Raden et al., 2018) was used to predict interactions between SviR and target transcripts. The −50 upstream sequence through stop codon of each PLE and ICP1 ORF was extracted. SviR-target predictions were calculated between extracted sequences and SviR using IntaRNA, requiring a maximum free energy of −10 (--outMaxE −10). For predictions of non-PLE 1 SviR alleles with target transcripts, SviR coding sequence was predicted based on homology to the PLE 1 allele. For mutational analysis of SviR-target basepairing, predictions were performed using CopomuS (Raden et al., 2020) using default settings.

### Plasmid construction

Genotypes of all plasmids are detailed in the strains and plasmids list. Oligonucleotides were purchased from Integrated DNA technologies. All plasmid constructs except CRISPR target constructs were generated through Gibson Assembly. Plasmids were linearized by PCR with Q5 polymerase using primers indicated in the oligonucleotides table. Inserts were amplified with Phusion polymerase using primers amplified with 20 nucleotide overhangs complimentary to the linearized vector. Gibson assembly was carried out with a 3:1 insert to vector molar ratio, with a total of 0.075 pmol DNA per reaction. 5 ul of the reaction was transformed in *E. coli* XL1-Blue chemically competent *E. coli* for further manipulation. pDD29 (T4 RNA ligase expression) was generated by linearization of p*tac*-RiboswitchE plasmid pKL06 and a codon-optimized *t4rnl1* gene insert.

Construction of CRISPR targeting constructs were generated through Golden Gate Assembly. Forward and reverse oligos complementary to the protospacer sequence with BsaI overhangs were mixed in 200 μL to a final concentration of 10 μM and boiled for 5 minutes to denature and cooled to room temperature. Oligos were phosphorylated with T4 Polynucleotide Kinase in 1x T4 DNA ligase buffer at 37°C for 30 min and heat inactivated for 20 minutes at 65°C. Reactions were dilution to 100 nM and Golden Gate Assembly reactions were carried out with BsaI restriction enzyme, 60 ng pCRISPR2.0 vector, and T4 DNA ligase at 37°C for 60 min.

### Bacterial strain construction

Generation of *V. cholerae* strains harboring plasmid constructs was carried out through conjugation with an *E. coli* S17 intermediate strain. Plasmid constructs were transformed into *E. coli* S17 and grown up overnight. Donor *E. coli* and recipient *V. cholerae* were mixed at an equal ratio of 0.5 ODs each, allowed to co-incubate for 1 hour, and plated on selective medium for recipient *V. cholerae* containing the plasmid of interest.

Generation of *V. cholerae* chromosomal mutants were generated with in-frame deletion of genes of interest using the allelic exchange vector pCVD442, encoding *sacB* enabling sucrose counter-selection. Template product was generated through splicing by overlap extension PCR to attach 1kb arms of up and downstream homology to the desired deletion and PCR products were inserted to the pCVD442 MCS by gibson assembly. Plasmids were mated into *V. cholerae* parental strains and exconjugants were grown overnight at 37°C in LB with ampicillin. Overnight cultures were plated on LB + 10% sucrose plates and incubated overnight at 30°C. Colonies were screened for loss of target deletions and confirmed by sanger sequencing.

### Phage mutant construction

Phage mutants were constructed as previously described (Box et al., 2016). *V. cholerae* expressing an inducible P*tac*:CRISPR-Cas system was transformed with pCRISPR2.0 plasmid containing spacer sequences targeting the stop codon of *gp120* and harboring a homologous repair template containing the desired mutation of interest. Cells were grown to OD_600_ of 0.2, induced with 1 mM IPTG, and infected with ICP1 following a 20 min pre-induction of Cas genes. Escape phages were isolated, passaged three times under CRISPR-Cas spacer only selection, and sanger sequenced to confirm desired mutation was acquired.

### RNA isolation

Bacterial strains were grown up to an OD_600_ of 0.3 and, when applicable, infected with phage at MOI=2.5. Samples were then collected at the desired timepoints, mixed 1:1 with ice-cold methanol if infected, pelleted by centrifugation, and snap-frozen in liquid nitrogen. Pellets were then processed following a standard TRIzol extraction protocol. Pellets were resuspended in 200 μL Tri Reagent (Millipore/Sigma) and incubated for 5 minutes at room temperature. Samples were mixed with 40 μL chloroform, mixed by inversion, and incubated for 10 min at room temperature. Samples were then centrifuged at 12,000xg for 10 min at 4°C. Following centrifugation, the upper (aqueous) phase was collected, promptly mixed with 110 μL 2-propanol and 11 μL pH 6.2 3M sodium acetate and mixed vigorously. Samples were centrifuged at 12,000xg for 15 min at 4° C, pellets were washed twice with 75% ethanol, and left to dry at room temperature. RNA was resuspended in 20-30 μL of DEPC water and RNA integrity and concentration were analyzed by NanoDrop.

### Hi-GRIL-Seq Sample Preparation

Hi-GRIL-seq was carried out based on previous protocols (Zhang et al., 2017), with some modifications. All Hi-GRIL-seq replicates were carried out using a *V. cholerae* PLE(+) host expressing T4 RNA ligase (DD25) during phage infection by ICP1_2011_AΔ*s9*. Cells were grown to an OD_600_ of 0.2 and induced with 1 mM IPTG and 40 mM theophylline for 20 minutes to pre-induce *t4rl1* expression. Induced hosts were infected at an MOI of 2.5 and allowed to progress through infection for 16 minutes, when 5 mL of cells were mixed 1:1 with ice cold methanol and RNA was isolated according the RNA isolation protocol above.

Following RNA isolation, each replicate was submitted to the UC Berkeley QB3 functional genomics lab, where ribosomal RNA was depleted according to the RiboZero rRNA Depletion kit (Illumina) and cDNA was synthesized. An S220 focused ultrasonicator (Covaris) was used to fragment the DNA, and library preparation was performed with the using the KAPA Hyper Prep kit for DNA (KK8504). Truncated universal stub adapters were ligated to each sample and adapter-ligated fragments were enriched for by PCR amplification with indexed primers. Samples were quantified with Quant-iT dsDNA Assay Kit (Life technologies), pooled equally by molarity, and submitted to the Vincent J. Coates Genomics Sequencing Laboratory at UC Berkeley, where libraries were sequenced on an Illumina HiSeq 4000 150 paired-end flow cell. RNA-seq data was analyzed as described under “quantification and statistical analysis”.

### 5’/3’-Rapid amplification of cDNA ends (RACE)

5’ and 3’ RACE were carried out on RNA isolated 16 minutes-post ICP1 infection of PLE(+) *V. cholerae,* as previously described (Argaman et al., 2001) with some modifications. 5’-RACE samples were prepared with 20 μg of RNA being treated with Tobacco Acid Pyrophosphatase (Millipore/Sigma) for 1 hour at 37°C to remove 5’ triphosphates. Reactions were quenched with Tri reagent and RNA was purified, with pellets resuspended in 20 μL DEPC water. RNA was dephosphorylated with Bacterial Alkaline Phosphatase (Thermo Fisher) in the presence of 20 U RNase inhibitor (Thermo Fisher) for one hour at 65°C to and reactions were inactivated with 1 μL of 0.5 M EDTA for 10 min at 50°C, and RNA was purified as described with Tri reagent. Pelleted, dephosphorylated RNA was resuspended in 10 μL DEPC water, mixed with 500 pmol of 5’ RNA adaptor A3, and annealed with T4 RNA ligase (NEB) overnight at 16° C. Ligated RNA was purified as described, resuspended in 37 μL DEPC water, and reverse transcribed with SuperScript III RT (Thermo Fisher) per manufacturers guidelines using a gene specific primer specific to the transcript of interest. First-strand cDNA was treated with RNAse H at 37°C for 20 min and subsequently amplified in a secondary reaction with primer A4 and a nested gene specific primer to amplify 5’ ends. Products were PCR cleaned up with Monarch PCR & DNA cleanup kit (NEB), ligated into pCR2.1-TOPO vector (Thermo fisher) per manufacturers guidelines, and transformed into XL1 Blue *E. coli.* At least 20 transformants were screened by colony PCR and 5’ TSS were identified from sanger sequencing of products.

3’-RACE was performed with 20 μg of equivalent RNA to 5’-RACE. RNA was dephosphorylated as described in 5’-RACE and purified as described with Tri reagent. Pelleted RNA was dissolved in 25 μL DEPC water, mixed with 500 pmol of RNA adapter E1, denatured for 5 min at 95° C, and chilled on ice for 5 min. 7 μL of RNA-adapter mix was mixed with 50 units of T4 RNA ligase, incubated overnight at 16°C, purified as described, and resuspended in 9 μL DEPC water. 1 μL of primer E4 was added to the ligated transcripts and cDNA synthesis was carried out with SuperScript III RT per manufacturers guidelines. cDNA was treated with RNAse H at 37°C for 20 min and a secondary PCR amplification was carried out with primer E4 and a gene specific primer, resulting in PCR amplicons containing 3’-RACE products. Ligation, transformation, colony PCR, and sequencing was performed as described for 5’-RACE, identifying 3’ ends of transcripts of interest.

### Northern blot analysis

For each sample, 5 μg of RNA was ran on a 8% polyacrylamide urea gel (National Diagnostics SequaGel 1:4) in 1x TBE buffer (100mM Tris base, 100m boric acid, 2 mM EDTA, pH 7.0) at 300V for 90 min. RNA was transferred from the gel to a Zeta-Probe GT membrane (Bio-Rad) for 16 h in 0.5X TBE at 20 V at 4°C. Transferred blots were UV-crosslinked on both sides and RiboRuler LR RNA Ladder (Thermo Fisher) was visualized and marked by UV illumination. Membranes were blocked for 2 h at 45°C with ULTRAhyb-Oligo Hybridization Buffer (Invitrogen). Northern probes were 5’ ^32^P-end labeled with 0.3 mCi of y-^32^P ATP (Perkin Elmer) with 10 U of T4 Polynucleotide kinase (NEB) for 1 h at 37°C. Labeled probes were spun through G-50 purification columns (Cytiva) to remove un-labelled probe, incubated at 95°C for 3 minutes, and added to blocked membrane. Membranes were hybridized for 16 h at 45°C in a rolling hybridization oven (Analytic Jena). Hybridized membranes were washed 2x with 20x SSC buffer (3M NaCl, 300 mM NaCit, pH 7.0), washed 3x with 2x SSC, and dried for 5 minutes at room temperature. Membranes were sealed in plastic binder pages and incubated with autoradiography film (Denville) and stored at −80°C for the appropriate exposure period, determined by probe signal.

### Western blot analysis

*V. cholerae* was infected with ICP1 at an MOI of 0.1 and grown until the indicated time point. 2 mL of infected cells were mixed with 2 mL ice cold methanol and centrifuged for 10 min at 5,000x g. Cell pellets were resuspended in ice cold lysis buffer (50 mM Tris, 150 mM NaCl, 1 mM EDTA, 0.5% Triton X-100, 1x Pierce Protease Inhibitor Mini Table (Thermo)). Protein concentration was calculated against a BSA standardized curve using with the Pierce BCA Protein Assay Kit and a total of 30 μg per sample was mixed with resuspended in 30 μL of 4x laemmli buffer (Bio-Rad) supplemented with 5% 2-mercaptoethanol and boiled for 10 minutes. 12 μL of sample was run on an Any-kD TGX-SDS-PAGE gel (Bio-Rad) in 1x TBS buffer for 30 minutes at 250 V and transferred using the Trans-Blot turbo Mini 0.2 μm PDVF transfer kit (Bio-Rad) with a Transblot Turbo transfer system (Bio-Rad) for 10 min at 2.5 V. Primary rabbit antiFLAG antibody (Sigma) was diluted 1:1500 and applied to membrane for 1 hour. Membrane was washed for 5 min 3x with TBS-T and incubated with secondary goat anti-rabbit-HRP antibody (Bio-Rad) at a 1:10000. Membrane was washed 3x for 5 min with TBS-T and developed with Clarity Western ECL (Bio-Rad) per manufacturers guidelines. Activated membranes were imaged using a Chemidoc XRS Imaging System (Bio-Rad).

### GFP reporter assay

GFP reporter assays were carried out as previously described (Corcoran et al., 2012). *E. coli* MG1655 was transformed with a *orf12.1::GFP* reporter plasmid and a *P_lac_-sviR* small RNA expression plasmid or empty vector control. Cells were grown up from single colonies overnight at 37°C and fresh cultures diluted to an OD_600_ of 0.05 in the presence of 1 mM IPTG. Cultures were grown for three hours and 1 mL culture was centrifuged at 5,000x g for 3 min and washed 1x with PBS. Washed pellets were resuspended in 2 mL 1x PBS and fluorescence was calculated using a LSR II cell analyzer (Beckman Coulter). Two biological replicates of 50,000 cell counts were analyzed with FlowJo, with mean fluorescent intensity of the GFP(+) population of cells reported for each replicate.

### Phage plaque assay

*V. cholerae* cells were grown to an OD_600_ of 0.2 and pre-induced for 20 minutes (if necessary). After 20 min pre-induction or at OD_600_ of 0.3 (without induction) 50 μL of culture was mixed with 10 μL of 10-fold serial dilutions of phage stocks. After 8 min incubation allowing for attachment of phages, infected cells were mixed with 4 mL 0.5% LB broth + 0.5% top agar in a 6 well tissue culture plates (Gennessee scientific) and incubated overnight at 37°C.

### Lysis kinetic assay

Lysis kinetic assays were carried out as previously described (Hays and Seed, 2019). *V. cholerae* cultures were grown to an OD_600_ of approximately 1.0 and back diluted to an OD_600_ of 0.05 before being allowed to grow to an OD_600_ 0.3. 150 μL of OD_600_ 0.3 was mixed with phage in a 96 well plate at an MOI of 2.5. Phage lysis was observed by measuring OD_600_ with an i3x SpectraMax plate reader at 37°C over 60 min with readings every 90 s, shaking in between reads.

### Genome replication qPCR

Genome replication assays were carried out as previous described (O’Hara et al., 2017). *V. cholerae* cultures were grown to an OD_600_ of approximately 1.0 and back diluted to an OD_600_ of 0.05 before being allowed to grow to an OD_600_ 0.3. 2 mL cultures were grown to an OD_600_ of O.3 at 37°C rolling and infected at an MOI of 0.1 (ICP1 replication) or 2.5 (PLE replication). 100 μL of culture volume was taken before (PLE replication) or immediately after (ICP1 replication) infection with ICP1, and boiled for 10 min. An endpoint sample was taken at 20 min post infection and boiled for 10 min. Boiled samples were diluted 1:50 (ICP1) or 1:1000 (PLE) in water and 3.75 μL of sample was mixed with 0.18 μL of 25 mg/mL primers and 7.5 μL of iQ SYBR master mix (Bio-Rad). 15 μL reactions were analyzed with a CFX connect real-time PCR detection system in technical duplicate and at least biological triplicate, and fold change was calculated by normalizing 20 min timepoints to input DNA calculated at 0 min timepoints.

