## Supplementary Figures and tables for "The RNA-RNA interactome between a phage and its satellite virus reveals a small RNA differentially regulates gene expression across both genomes"

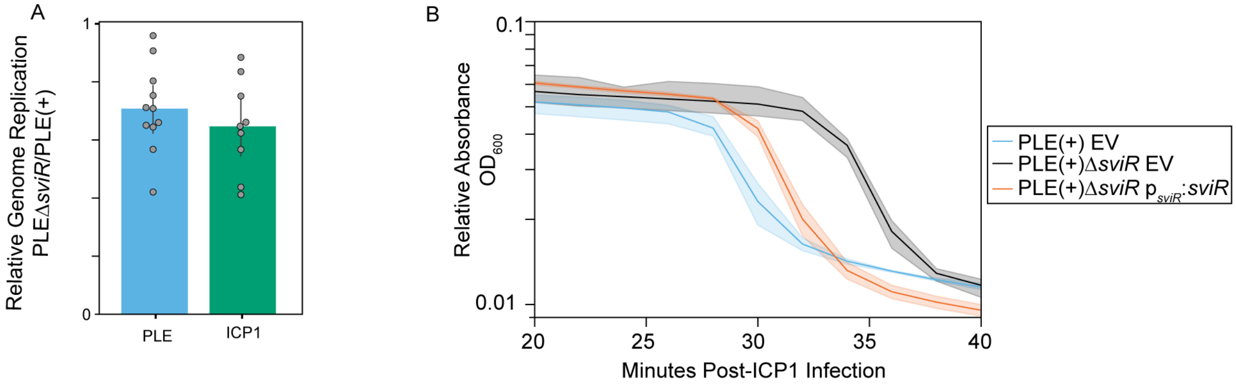


Figure S1 (Supplemental to Figure 1) SviR is necessary for wild type PLE response to ICP1 infection
(A) Genome replication of both PLE and ICP1 are decreased in the absence of SviR. qPCR was used to measure genome replication of both PLE and ICP1 during the ICP1 infection cycle. Relative genome replication is measured as the fold from 0 to 20 minutes in a PLEΔ*sviR* cells normalized to the fold change in wild type PLE(+) cells of paired samples.
(B) Timing of *V. cholerae* lysis kinetics of PLEΔ*sviR* cells is delayed in compared to PLE(+) cells. Strains were infected with ICP1 at a high MOI and lysis kinetics of infected cells were surveyed over time. Solid lines represent average of three replicates and shaded outlines represent ± standard deviation of replicates.

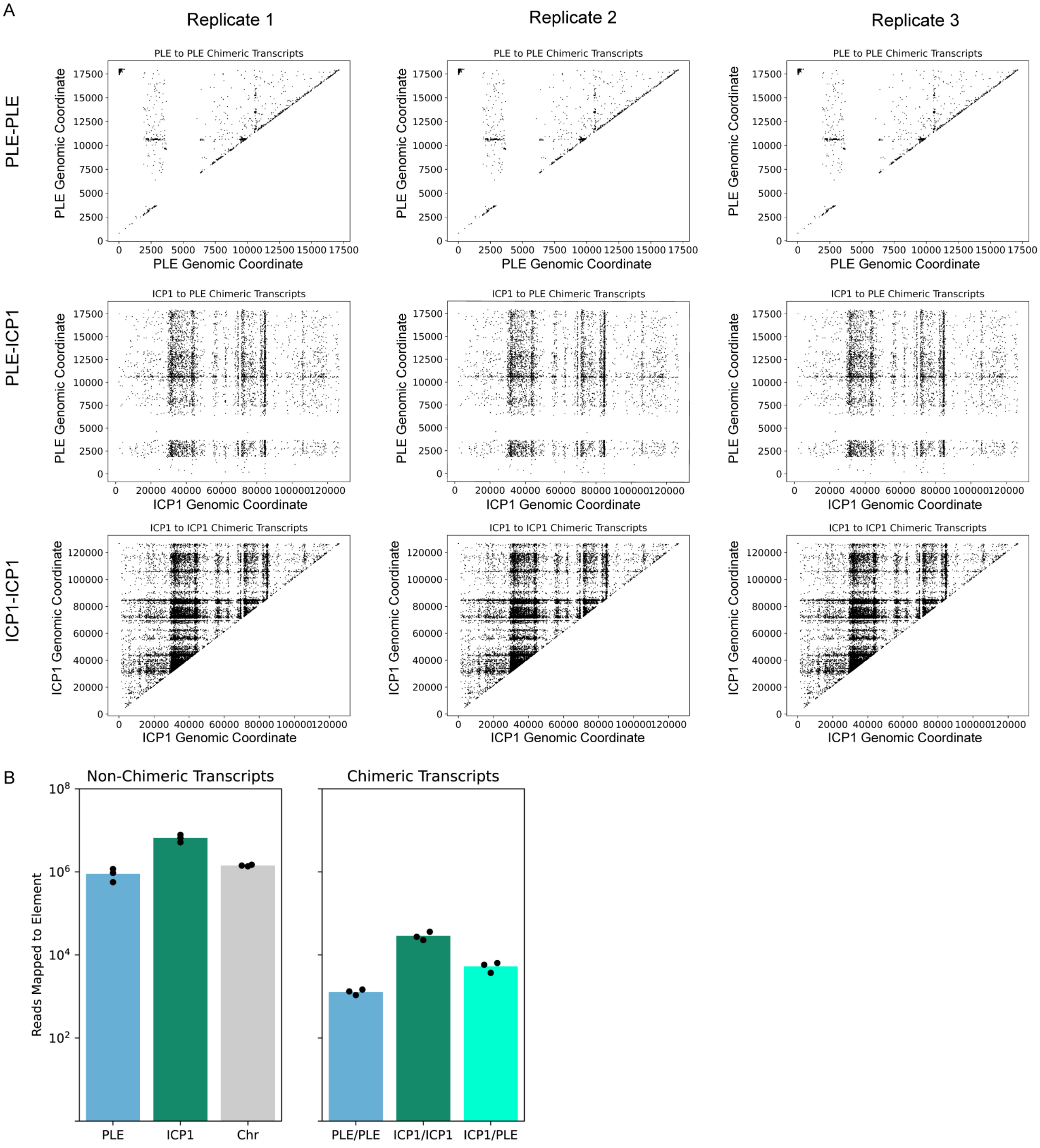


Figure S2 (Supplemental to Figure 2) End mapping of chimeric reads between ICP1 and PLE between biological replicates

(A) Mapping of X-Y coordinate pairs for the ends of each Hi-GRIL-seq chimeric read, as performed for figure 2. Each column represents reads from one replicate and with each row representing a reference genome pairing. Read locations were determined as described in figure 2B-D and star methods.

(B) Proportion of reads between each biological replicate mapping to each genome/genome pairing. Inter-genome chimeric transcripts are separated by at least 1000 bp, comparable to other Hi-GRIL-seq analyses.

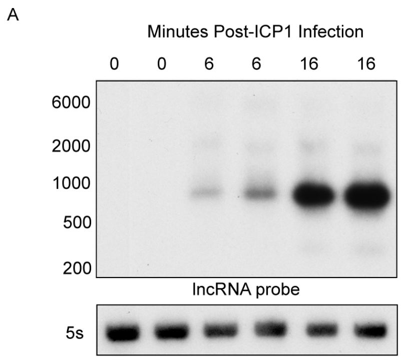


Figure S3 - Supplemental to Figure 2) Northern blot of the ICP1 encoded lncRNA over a time course of ICP1 infection of PLE(+) *V. cholerae*

RNA was isolated from two biological replicates of a high MOI ICP1 infection of PLE(+) at 0, 6, and 16 minutes post-ICP1 infection and isolated RNA was probed for the ICP1 lncRNA. Expression of the lncRNA increases throughout infection, comparable to that of SviR

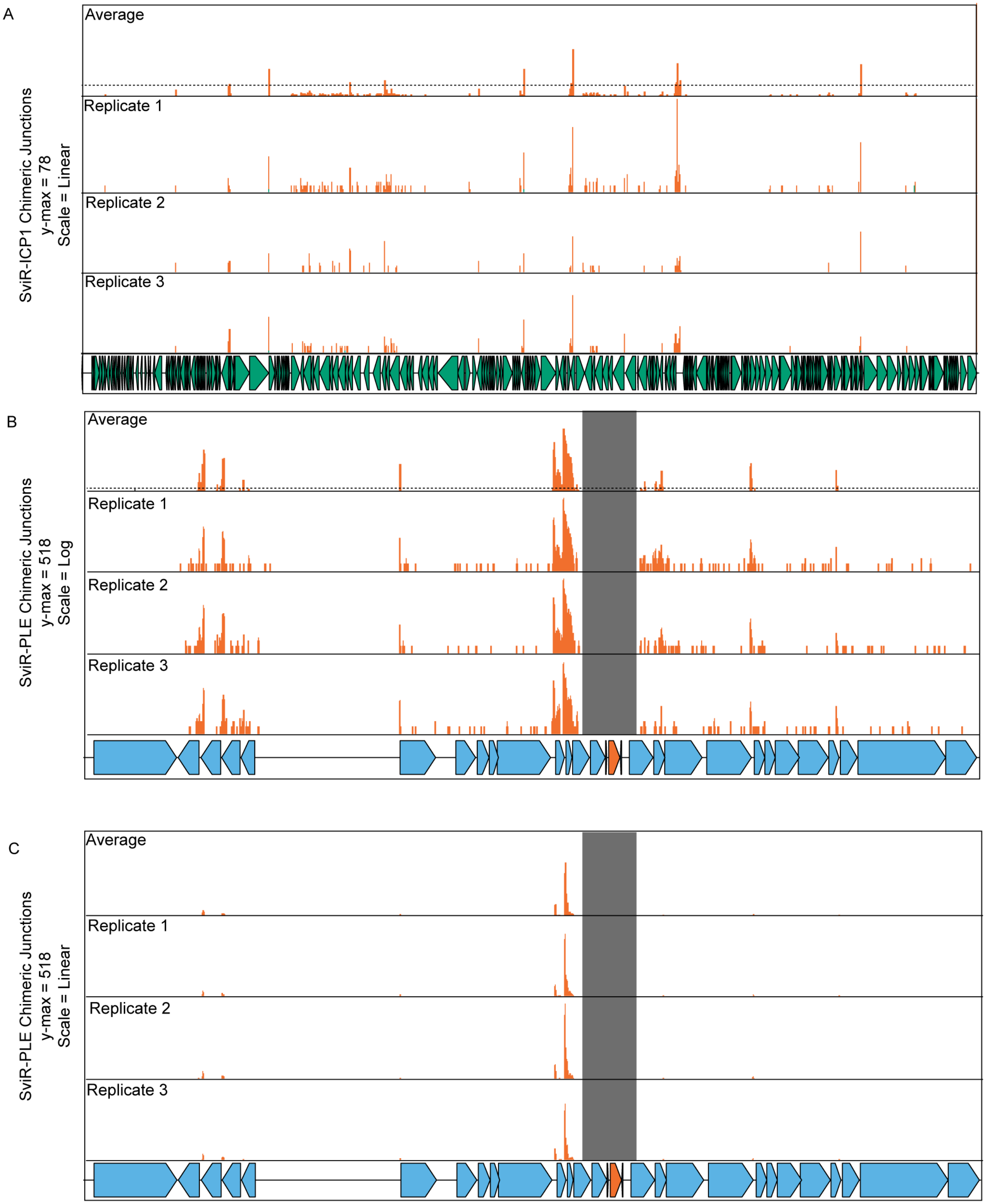
A
SviR-ICP1 Chimeric Junctions
y-max = 78
Scale = Linear
Average
Replicate 1
Replicate 2
Replicate 3
B
SviR-PLE Chimeric Junctions
y-max = 518
Scale = Log
Average
Replicate 1
Replicate 2
Replicate 3
C
SviR-PLE Chimeric Junctions
y-max = 518
Scale = Linear
Average
Replicate 1
Replicate 2
Replicate 3

Figure S4 (Supplemental to Figure 4 and 5) Hi-GRIL-Seq junction mapping and filtering of SviR-target chimeras to PLE and ICP1

(A, B, and C) Mapping of SviR-target chimeras to the ICP1 genome (A) and PLE (B and C) genomes. The 25 target base pairs adjacent to the SviR-target junction were identified as described in Figure 4,5, and star methods, and mapped to reference genomes. Individual replicates have all reads mapped (below), whereas averaged samples (top) only show positions above the noise threshold of 10 or more reads consecutively mapping to a single locus (dashed line). Gene graphs below diagrams represent ICP1 (green) and PLE (blue) ORFs, and the SviR (orange). Scales and y-max values are indicated on the y axis of each replicate (A and C = linear, B = log). Black shaded box (B and C) indicate the region within 1 kb from the center of SviR, where chimeric reads were unable to be discerned from transcripts.

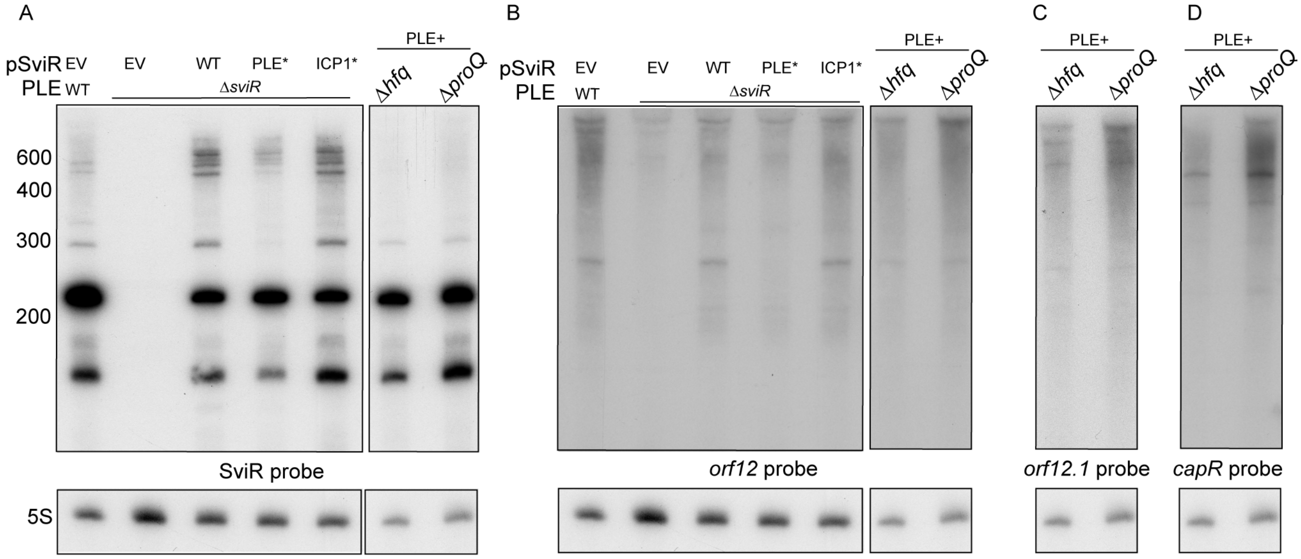


Figure S5 (Supplemental to Figure 5) SviR allele expression and regulation without RNA chaperones
(A) SviR mutant allele expression (left) and wild type chromosomal expression in the absence of RNA chaperones Hfq and ProQ (right). SviR mutant alleles are expressed relatively equally from pSviR plasmids. Chromosomal expression of SviR is comparable in the absence of RNA chaperone Hfq, although expression levels may be slightly decreased.
(B) Northern blot analysis of PLE *orf12* regulation by SviR occurs dependent on the PLE seed region and independent of RNA chaperones Hfq or ProQ.
(C and D) Regulation of *orf12.1* and *capR* occurs independent of RNA chaperones Hfq or ProQ.
Duplicated 5S blots between A-D are present due to stripping and re-probing of the same blots.

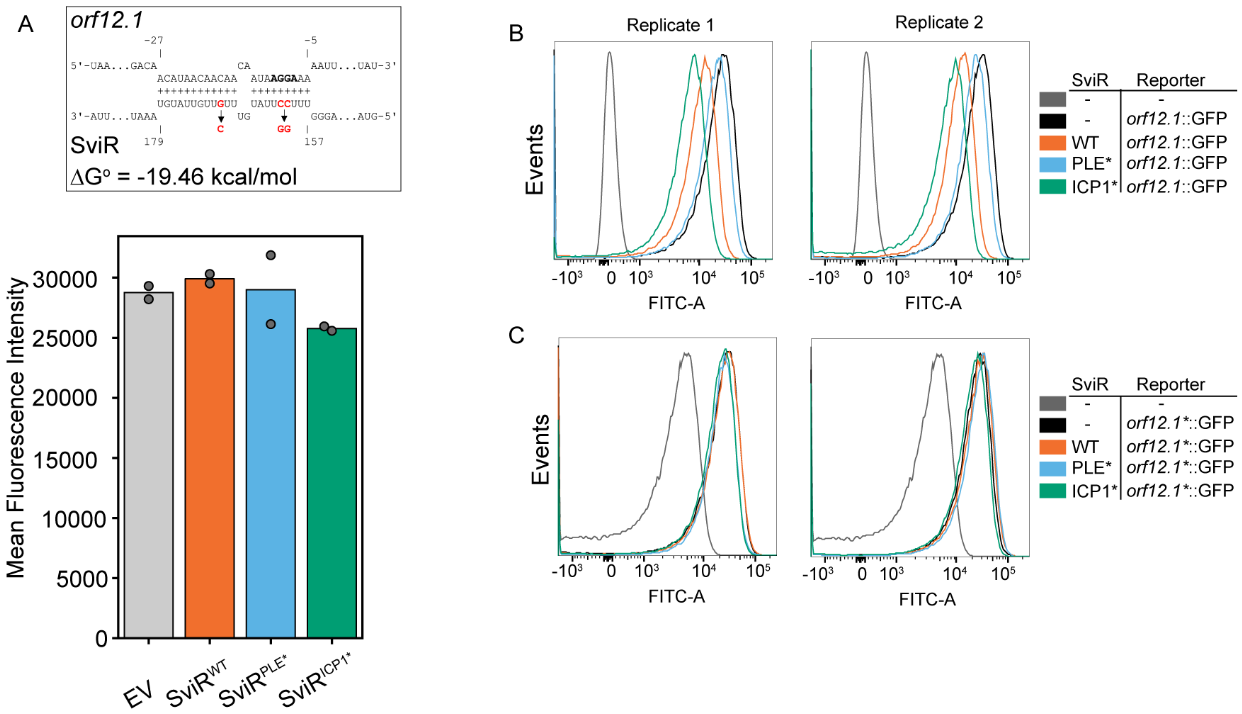


Figure S6 – (Supplementary to Figure 5) Mean fluorescent intensity of *orf12.1**::GFP, containing compensatory mutations for interactions with SviR$^{PLE*}$, is inconclusive.

(A) Mutations to the RBS of *orf12.1* (IntaRNA, red bases) were made to generate *orf12.1**::GFP compensatory mutation reporters. Average mean fluorescence intensity (MFI) from 50,000 events is reported on the y-axis.

(B and C) Histograms generated with FlowJo representing the fluorescence intensity of total populations of cells with various reporter constructs and SviR alleles. Cells were grown for three hours and resuspended in 1X PBS, then subjected to flow cytometric analysis. Photomultiplier tube voltage parameters vary considerably between B (FITC = 643 V) and C (FITC = 950 V) making values of FITC histograms and MFI between the two experiments incomparable.

| Target Transcript | SviR Range | Predicted Interaction $\Delta G^o$ | Target Range |
|---|---|---|---|
| *ORF4* | 158 to 177 | -12.75 kcal/mol | -9 to -29 |
| *ORF5* | 157 to 174 | -14.24 kcal/mol | -31 to -11 |
| ***ORF13*** | 154 to 174 | -10.79 kcal/mol | -24 to +8 |
| *nixI* ***(ORF15)*** | 156 to 177 | -11.70 kcal/mol | -7 to -27 |
| ***ORF16*** | 156 to 175 | -17.14 kcal/mol | -7 to -26 |
| *tcaP (ORF17)* | 157 to 179 | -18.93 kcal/mol | -9 to -39 |

**Table S1 (Supplementary to table 1 and figure 5) IntaRNA predictions for non-Hi-GRIL-Seq PLE Targets**

IntaRNA predictions between SviR and PLE ORFs that were not observed as chimeras in Hi-GRIL-Seq above the detection threshold. The interactions are all between the SviR PLE seed region experimentally validated in Figure 5 and the 5’ UTR of ORFs. SviR range and target range indicate the base pairs predicted to interact by IntaRNA. Stability of these predicted interactions is represented by the prediction interaction $\Delta G^o$ column. Putative targets highlighted in bold represent transcripts within the 1kb cutoff from the center of SviR, and as such Hi-GRIL-seq interactions were unable to be detected for these samples
